# OKN4395, a first-in-class EP2/EP4/DP1 triple antagonist reprograms prostanoid-driven immunosuppression to restore antitumor immunity

**DOI:** 10.64898/2026.02.08.704632

**Authors:** Maximilien Grandclaudon, Morgane Boulch, Anna Thaller, Jonathan Sabio Ortiz, Alexandre Grimaldi, Marie Goxe, Agnès Knopf, Marie Daugan, Eva Huehn, Carmela Gnerre, Sébastien Jeay, Monica Faronato, Haithem Dakhli, Silvia Lopez-Lastra, Alexandra Hardy, Sandrine Sanchez, Imke Mayer, Rose Hoste, Floriane Montanari, Vassili Soumelis, Joan Alberti, Lucia Pattarini, Caroline Hoffmann, Andrew J. Pierce

**Affiliations:** Owkin; Idorsia Pharmaceuticals Ltd; Idordia Pharmaceuticals Ltd; Admetra DMPK/PD consultancy

## Abstract

Immune checkpoint inhibitors, particularly T cell targeting anti-PD(L)1 therapies, have revolutionized the treatment landscape for solid malignancies, but challenges related to non-responsiveness and the development of treatment resistance continue to be observed. An additional immunosuppressive axis relates to prostaglandin signaling downstream of cyclooxygenase-2 (COX2), where COX2 inhibitors have shown clinical promise in re-engaging both T and non-T cell immune compartments, yet have suffered from toxicity concerns. We report here the preclinical characterization of OKN4395, a highly potent and specific first-in-class triple antagonist of EP2, EP4, and DP1, major tumor immunosuppressive receptors downstream of COX2. OKN4395 restores immune function on both T cells and NK cells *in vitro*. Additionally, OKN4395 acts synergistically with anti-PD1 to increase speed and depth of antitumor activity. Overall, these findings robustly support the clinical investigation of OKN4395 in an ongoing Phase 1 trial (NCT06789172) as an innovative cancer immunotherapy for solid tumors, as a single agent and in combination with anti-PD1 therapy.

**Statement of significance:** OKN4395, a first-in-class oral EP2/EP4/DP1 antagonist, reverses prostanoid-driven immunosuppression to restore antitumor immunity. Integrated pharmacology defines mechanism, translational biomarkers as well as both monotherapy and anti-PD1 combination strategies. These data position prostanoid tri-receptor antagonism as a translatable strategy in solid tumors. A global Phase 1 study is underway (NCT06789172).

## INTRODUCTION

The arachidonic acid (AA) metabolic pathway has long been associated with cancer progression^1^. Within this pathway, prostaglandin E2 (PGE2), a critical inflammatory lipid produced by cyclooxygenase-2 (COX2, HGNC: 9605), its rate limiting enzyme, has been identified as a key mediator of AA pathway activity in cancer. Substantial concentrations of PGE2 have been reported in various human tumors^2^. The pro-tumoral functions of PGE2 have been mechanistically well-characterized in preclinical oncology models, affecting both tumor cell growth and various components of the tumor microenvironment (TME)^3^. Specifically, pronounced inhibitory effects have been demonstrated for PGE2 on the immune compartment, including T cells (both CD4^+^ and CD8^+^)^4^ and natural killer (NK) cells^5,6^, preventing their cytotoxic effects towards the tumor compartment. Two recent studies shed light on downregulation of IL-2R on T cells as a critical mechanistic event occurring downstream of PGE2 within the TME and revealing one potent and long-lasting cellular effect for PGE2^2,7^. In addition, PGE2 is also known to impact myeloid cell biology by promoting the polarization of anti-inflammatory tumor-associated macrophages (TAMs) ^8,9^ and inhibiting maturation of dendritic cells (DCs)^10^. Recent studies further highlighted the importance of PGE2 in promoting IL-1β producing macrophages in a murine pancreatic ductal adenocarcinoma (PDAC) model matching a subset of macrophages observed in human settings^11^. Moreover, others demonstrated a causal relationship between MAPK signalling in cancer cells and enhanced PGE2 secretion, which explained the absence of strong inflammatory monocyte infiltration and active intratumoral T cell antitumoral responses^12^. Collectively, these preclinical insights reveal deep implication of the PGE2 pathway in hindering the cancer immunity cycle through direct impacts on both innate and adaptive mechanisms.

These findings collectively suggest that the COX2-PGE2 axis is central and potentially dominant in specific tumor contexts. Blocking this pathway using non-steroidal anti-inflammatory drugs (NSAIDs) or COX2 inhibitors has demonstrated potential benefits across various solid tumor indications^13–18^. Clinical trials combining COX2 inhibitors (e.g., celecoxib) with anti-PD1 therapies have also shown promise^19^.

However, the clinical use of these approaches has been limited due to a lack of specificity and toxicity. Blocking COX or COX2 affects multiple prostaglandin pathways, which can lead to dose-limiting cardiac or renal toxicities^20^. Recent research suggests that specifically blocking the production of cyclic AMP (cAMP) modulated by PGE2 downstream of its receptors may promote antitumor immunity while minimizing toxicity. Based on these findings and in order to cope with the potential side effects of COX2 inhibition, several groups developed prostaglandin E receptor 4 (EP4) specific inhibitors that are currently under active clinical development^21–24^. These strategies have shown potential and signs of clinical activity in various contexts, particularly in colorectal cancer (CRC) and gastric cancer^25^.

Importantly, several mechanistic insights revealed that PGE2 could signal via both prostaglandin E receptor 2 (EP2) and EP4 within the TME in a redundant fashion, highlighting that EP4 blockade alone could be compensated by PGE2 signaling via EP2^26,27^. Several preclinical studies confirmed that full efficacy of PGE2 inhibition signaling should therefore be reached by dual EP2/EP4 inhibition^28^.

In addition, while investigating a human melanoma cohort resistant to anti-PD1 therapy, a recent study identified that prostaglandin D2 (PGD2), another lipid mediator downstream of COX2, could play an important immunosuppressive role through the DP1 receptor signaling in immune cells^29^. Specifically, the authors highlighted that PGD2 could exhibit dual modes of action in the TME. Firstly, it can function in an autocrine fashion, enhancing the anti-inflammatory phenotype of macrophages. Secondly, PGD2 can operate in a paracrine manner, activating DP1 signaling pathways in CD8^+^ T cells resulting in the inhibition of CD8^+^ T cell recruitment, activation, proliferation, and antitumor cytotoxic activities. Importantly, both PGE2 and PGD2 suppress immune effector function through a common mechanism: elevation of intracellular cAMP levels in T and NK cells via their respective Gs-coupled receptors (EP2/EP4 and DP1)^27,30^. Because these pathways utilize the same second messenger in the same target cells, they represent functionally redundant immunosuppressive signals, raising concerns about potential cross-resistance when inhibiting either pathway in isolation.

In this study, we provide a comprehensive preclinical pharmacology overview of OKN4395, to our knowledge a first-in-class triple specific small molecule antagonist of EP2, EP4 and DP1 prostanoid receptors for solid tumor treatment. This approach aims to strike a balance between strategies that may be either too broad or too narrow: upstream COX2 inhibition, while effective, disrupts multiple prostaglandin pathways leading to toxicities that have limited achieving full therapeutic benefit in oncology; conversely, highly selective single-receptor antagonists such as EP4 inhibitors may be insufficient given the mechanistic redundancy of cAMP-mediated immunosuppression through EP2 and DP1. By simultaneously blocking all three major prostanoid receptors that converge on cAMP signaling in immune effector cells, OKN4395 is designed to maximize blockade of prostaglandin-driven immunosuppression while maintaining specificity to avoid the broader toxicities associated with upstream COX2 inhibition.

## RESULTS

### OKN4395 is a selective and equipotent triple antagonist of EP2, EP4 and DP1 receptors

OKN4395 originates from a prostanoid-receptor antagonist series informed by prior EP2/EP4 medicinal chemistry^31–33^. Guided by shared pharmacophore features reported for EP2/EP4 antagonists, we optimized a di-aryl core bearing an anionic functionality and heteroatom-containing linker elements to support receptor engagement.

First, the activity of OKN4395 was determined using a functional cAMP assay on HEK293 cells stably transfected for either human EP2 or human EP4 receptors treated with PGE2. For this purpose, PGE2 was titrated and used at its EC80 concentration. In this assay, OKN4395 was found to exhibit potent dual antagonistic activity against the human PGE2 receptors, EP2 and EP4. The compound demonstrated to be of high potency, with mean IC50 values of 8.1 nM for EP2 (Figure 1B) and 26.6 nM for EP4 (Figure 1C) in these cellular assays. Similar values and inhibition patterns were found when assessing murine, rat, cynomolgus and dog EP2 and EP4 orthologues (Figure S1A). Challenging OKN4395 with increasing concentrations of PGE2 using the SF295 and BT549 cell lines naturally expressing either human EP2 or EP4 revealed the surmountable nature of the OKN4395 antagonist (Figure S1B). Taken together, these results indicate that OKN4395 is a highly effective dual antagonist of both EP2 and EP4 receptors.

**Figure 1:**
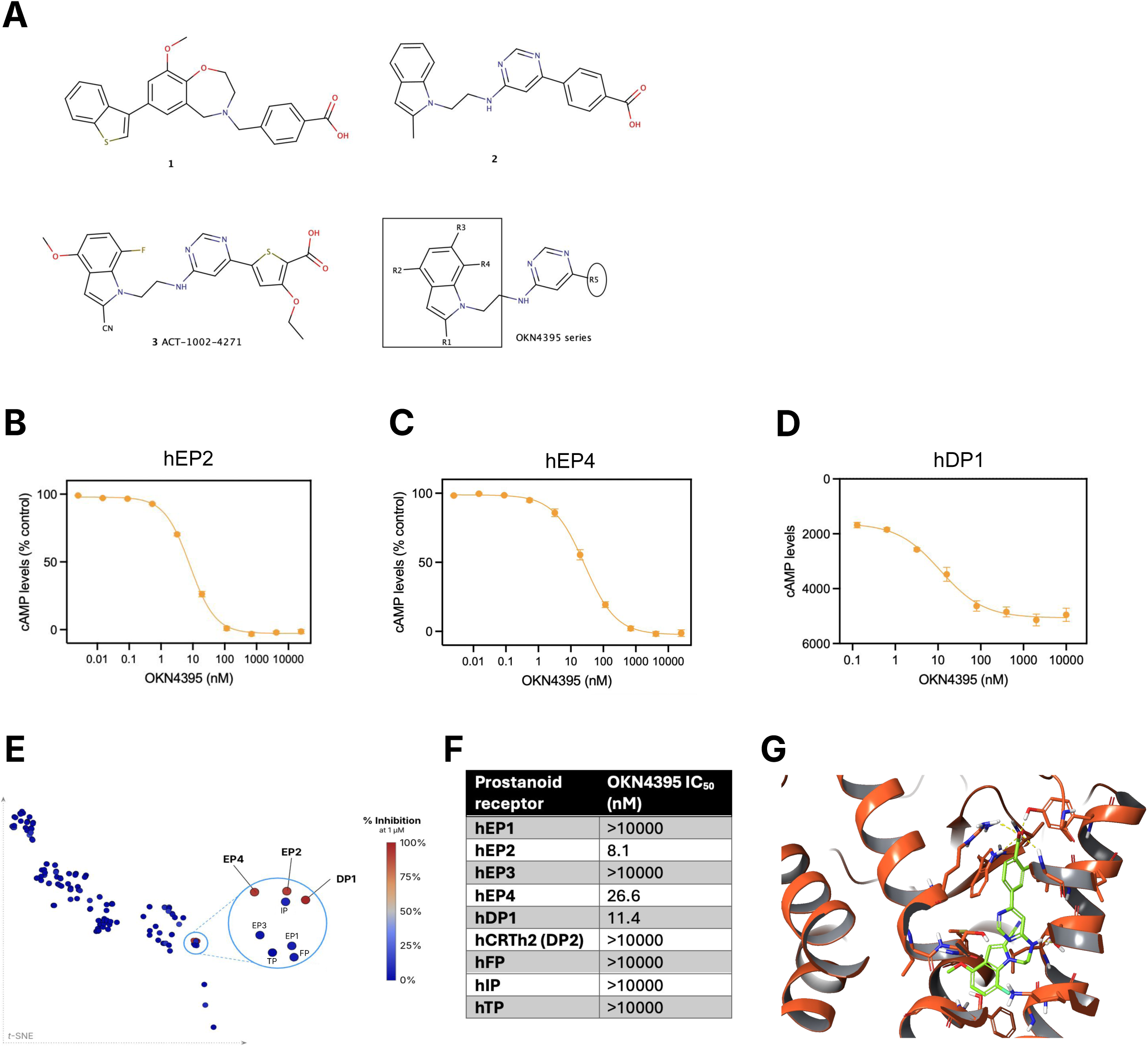
OKN4395 is a potent and selective inhibitor of EP2, EP4 and DP1. (**A**) Compound 1: lead compound from [29] with moderate dual inhibition; Compound 2. lead compound from [30] with potent EP2 inhibition; Compound 3. lead compound from [31] with potent dual EP2/EP4 inhibition. The OKN4395 series is derived from [31]. Most of the diversity in the series comes from the exploration of the right hand aryl system bearing the carboxylic acid. Compound 3 is an example of that series. (**B**) cAMP production after stimulation with PGE2 (EC80, 2nM) measured on HEK293 stable clones expressing human EP2 (hEP2). Values represent mean of n=10 technical replicates ± SEM. (**C**) cAMP production after stimulation with PGE2 (EC80, 6nM) measured on HEK293 stable clones expressing human EP4 (hEP4). Values represent mean of n=10 technical replicates ± SEM. (**D**) cAMP production after stimulation with PGD2 (EC80, 300pM) measured on CHO-K1 cells stable clone transfected for human DP1 (hDP1). Values represent mean of n=4 technical replicates ± SEM. (**E**) Visualization of OKN4395 antagonist activity at 1µM screened with gpcrMAX GPCR LeadHunter Panel (Eurofins) obtained with the plotrender spatial data mapper tool developed by gpcrdb. (**F**) Summary table of OKN4395 IC50 on human prostanoid receptors. (**G**) Plausible binding mode of a representative of OKN4395 series in DP1 (DP1: orange, ligand: green). Hydrogen bonds are yellow dashed lines.

Given the potential relevance of the PGD2-DP1 axis, we wanted to carefully assess the potential activity of OKN4395 on cAMP induced by DP1 signaling. Using CHO-K1 cells stably transfected for the human DP1 receptor, full titration of OKN4395 demonstrated a very potent inhibition of cAMP signaling induced by PGD2, with mean IC50 value of 11.4 nM (Figure 1D). To validate specificity of action of PGD2 on DP1, we validated that parental CHO-K1 did not induce cAMP response in similar PGD2 stimulation (data not shown).

We then addressed OKN4395 selectivity towards closely structurally related prostanoid receptors. Using a similar methodology than for EP2, EP4 and DP1 potency studies, we showed that OKN4395 had no activity on human EP3, human EP1 and human IP receptors even when OKN4395 was used at very high concentrations (>10µM) (Figure S1C). To further understand whether OKN4395 was reactive against other members of the GPCR family, a comprehensive screen using 1µM or 10µM of OKN4395 was performed across 166 GPCR searching for both agonistic or antagonistic activities. This large screen revealed that no GPCR receptor other than EP2, EP4 and DP1 displayed more than 20% of activity at the 1µM dose (Figure 1E). Specifically, this large test also included FP, TP and CRTH2 (DP2), three prostanoid receptors structurally similar to EP2, EP4 or DP1. Absence of OKN4395 activity was demonstrated for FP, TP and DP2, even for the highest dose used of 10µM (Figure 1F, Figure S1D).

Finally, we wanted to understand the binding mode of OKN4395 and its series to EP2, EP4 and DP1 and how the structure of the proteins could explain the triple activity. To do so, docking experiments were performed for the three proteins. This analysis revealed a network of interactions extremely similar between the three targets (Figure 1G, Figure S1E): 1/ Stabilization of the negative charge of the carboxylic acid by an arginine and at least two sidechains containing hydrogen-bond donors such as a tyrosine, a threonine and a tryptophan. 2/ Anchoring of the aminopyrimidine moiety through a hydrogen bond with a serine residue localized two turns before the above-mentioned tyrosine in the same alpha helix (i, i+7 positions). Both of these clusters of interactions were found to be stable in molecular dynamics run in EP4 and DP1. Superimposition of the three binding sites revealed that this interaction system is the same across the three proteins, as the key residues are conserved throughout (Figure S1E). To further highlight the importance of this interaction network, we studied the binding site of EP3 in which the serine is replaced by a threonine. This simple change blocks the hydrogen bond formation due to the added methyl group as seen in molecular dynamics run analysis (Figure S1F) and is thought to be a cause of the inactivity observed against EP3. As was observed both visually and through plotting relevant distances (hydroxy group to nitrogens of the aminopyrimidine), no strong hydrogen bonds were established, resulting in a global shift of the binding pose. This shift weakened the salt-bridge established between the carboxylate and the arginine residue found in all four proteins. Overall, these results reveal the unique profile of OKN4395 as a triple equipotent and specific antagonist—targeting only EP2, EP4 and DP1 while sparing other prostanoid receptors and the broader GPCR family.

### OKN4395 restores PGE2-mediated suppression of both T and NK cell activities

OKN4395 was subsequently studied in a series of functional assays using primary human immune cells to characterize its mode of action. Mirroring what was performed in cell line models, we studied OKN4395 inhibition of PGE2 induced cAMP in total CD3^+^ human primary T cells, models closer to human physiopathological settings. Using RT-qPCR, expression of EP2 and EP4 receptors was confirmed in primary T cells. In addition, EP1 and EP3 receptors were not detectable, revealing that these two receptors are unlikely to mediate PGE2 signaling in our assay (Figure S2A). In order to use a relevant PGE2 concentration, we titrated its effect and determined the EC90 for cAMP induction in T cells (Figure S2B). At EC90 concentration of PGE2, we were able to demonstrate that PGE2 induced cAMP production was fully blocked only when combining both EP4 and EP2 inhibition. In this system, inhibition of EP4 alone was fully compensated by PGE2 signaling through EP2, demonstrating that dual inhibition is a prerequisite to totally block PGE2 downstream and restore T cell activation. In this assay, OKN4395 demonstrated higher potency than combination of EP2 and EP4 single inhibitors and reached similar efficacy levels highlighting its functional dual EP2/EP4 inhibition capacity (Figure 2A). Using an equivalent methodology but replacing human T cells by human primary NK cells, we first confirmed the functional expression of EP2 and EP4 receptors by NK cells (Figure S2A) and PGE2-mediated cAMP signaling in this cell type (Figure S2C). Then, we similarly demonstrated that OKN4395 mediates a strong inhibition of the cAMP induction by PGE2 in this distinct effector immune cell population (Figure 2B).

**Figure 2:**
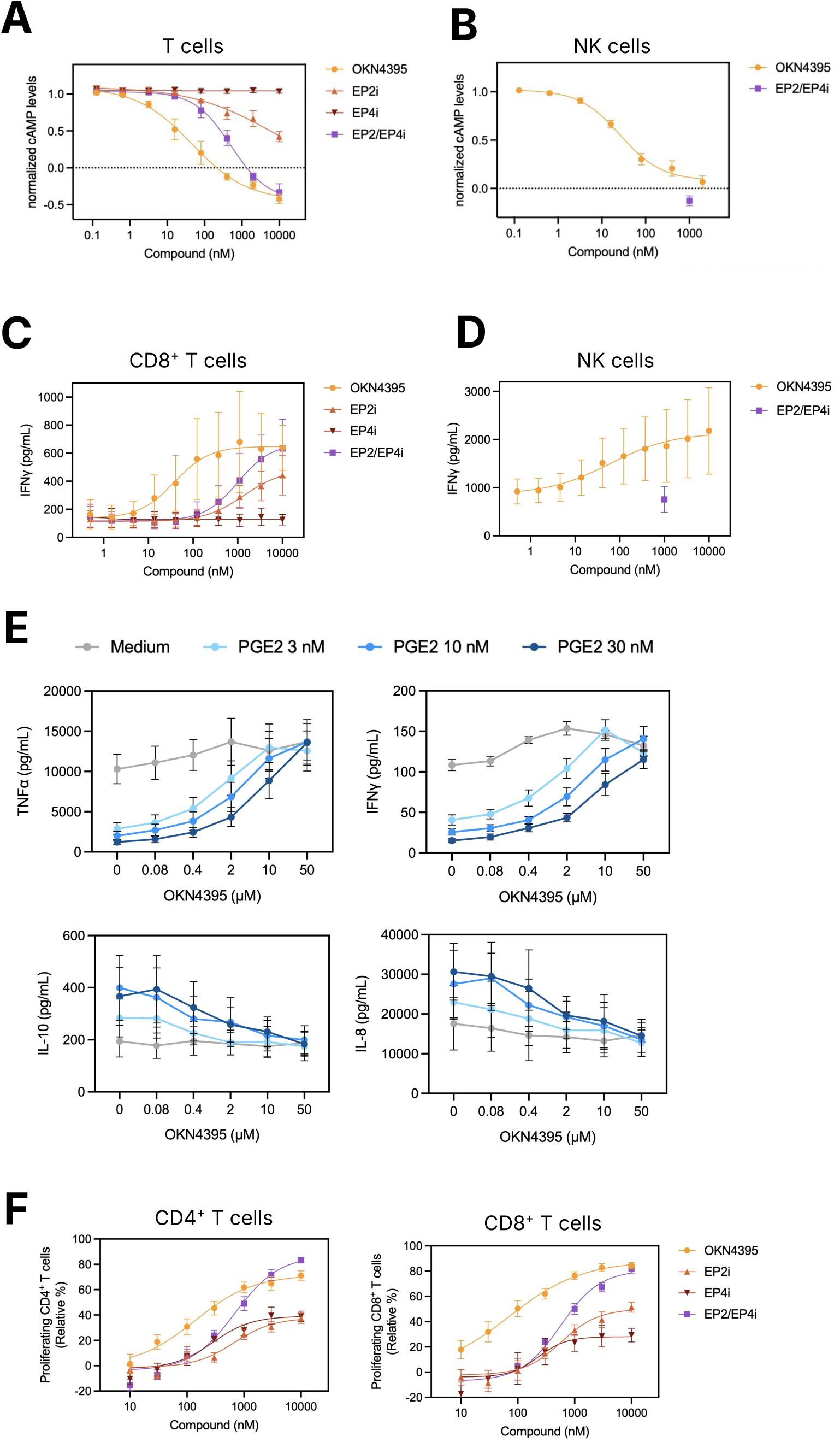
OKN4395 Alleviates PGE2-Induced Suppression of Immune Cell Functions In Vitro. (**A**) cAMP production of Pan T cells in the presence of PGE2 (EC90, 7.8nM). Values represent mean of n=3 healthy donors ± SEM. (**B**) cAMP production of NK cells in the presence of PGE2 (EC90, 30nM). Values represent mean of n=3 healthy donors ± SEM. (**C**) IFNγ secreted by CD8^+^ T cells stimulated with CD3/CD28 Dynabeads in the presence of PGE2 (200nM). Values represent mean of n=3 healthy donors ± SEM. (**D**) IFNγ secreted by NK cells stimulated with IL-2, IL-12, IL-15 and IL-18 in the presence of PGE2 (1000nM). Values represent mean of n=3 healthy donors ± SEM. (**E**) TNFα, IFNγ, IL-10 and IL-8 concentrations recovered in the supernatant of a human whole blood test engagement in the presence of LPS (0.1µg/mL) and increasing concentrations of PGE2 (0-3-10-30nM) for 24 hours. Values represent mean of n=3 healthy donors ± SEM. (**F**) Proliferation of CD4^+^ (left) and CD8^+^ (right) T cells stimulated with CD3/CD28 Dynabeads in the presence of PGE2 (500nM). Values represent mean of n=4 healthy donors ± SEM.

Since these results are of importance regarding OKN4395 upstream cAMP regulatory functions in a PGE2 context, we wanted to further investigate OKN4395 pharmacology in assays monitoring established downstream immune cell functions. First, we set up an *in vitro* assay measuring IFNγ production by purified human CD8^+^ T cells following activation with anti-CD3/CD28 activation beads. In this assay, PGE2 was able to fully inhibit IFNγ secretion (Figure S2D). OKN4395 restored IFNγ production in a dose-dependent manner following a similar pattern than for the cAMP readout. Using specific EP2 and EP4 inhibitors as controls, we were also able to demonstrate that inhibition of both receptors was needed to reach maximal efficacy (Figure 2C). Similarly, PGE2 significantly inhibited IFNγ production from activated human NK cells *in vitro* (Figure S2E) and OKN4395 was able to completely restore their IFNγ secretion (Figure 2D).

To further characterize the impact of OKN4395 on cytokine and chemokine secretion, we studied its ability to suppress PGE2-mediated effect in an *ex vivo* whole blood assay, where total blood was stimulated by LPS during 24 hours at 37°C in presence of distinct concentrations of OKN4395. At higher concentrations, OKN4395 exhibited remarkable potency in fully restoring the co-inhibited cytokine responses. This effect was most prominent for TNFα and IFNγ (Figure 2E). IL-6 and CCL3 also showed significant restoration (Figure S2F). Interestingly, PGE2 actually enhanced the release of IL-10 and IL-8, two cytokines renowned for their inhibitory and pro-tumoral properties. In this unique scenario, OKN4395 did not further elevate IL-10 or IL-8 concentrations in the supernatants but instead induced a dose-dependent decrease (Figure 2E).

Finally, using an *in vitro* proliferation assay, we determined the impact of PGE2 treatment on activated CD4^+^ and CD8^+^ T cells and studied functional blockade using OKN4395. In this assay, we were able to determine that both EP2 and EP4 single inhibitors when used separately were able to only minimally restore both CD4^+^ or CD8^+^ T cell proliferation. However, once both inhibitors were combined, or when using OKN4395, proliferation of both CD4^+^ and CD8^+^ T cells was dramatically impacted and fully restored (Figure 2F).

### PGD2-mediated immunosuppression is fully blocked by OKN4395

Then, we studied more closely OKN4395-mediated inhibitory activity of DP1 using a series of PGD2-dependent *in vitro* primary immune cell assays. First, we determined if, similarly to PGE2, PGD2 was able to induce cAMP in total CD3^+^ primary human T cells. We demonstrated that T cells express substantial levels of DP1 receptor while DP2 receptor was barely expressed (Figure S2A). Interestingly, we retrieved a strong induction of cAMP by PGD2 with equivalent levels to what was seen for PGE2 (Figure S3A). Using high concentrations of PGD2 (EC90, 92nM), we demonstrated that OKN4395 was able to reduce in a dose-dependent manner the measured cAMP induction to similar levels than laropiprant, a well known extremely potent DP1 inhibitor (Figure 3A). We also assessed the PGD2 specificity of signaling through DP1 in this assay, by showing complete absence of effect of EP2 and EP4 combined inhibitors. Using primary NK cells, we observed a similar impact of PGD2 on cAMP levels than for primary T cells (Figure S3B) and OKN4395 was also fully able to inhibit PGD2 induction of cAMP in NK cells (Figure 3B), consistent with their DP1 expression (Figure S2A). As shown for PGE2, the PGD2 impact on upstream cAMP regulation also dramatically impacted IFNγ secretion capacity for both T cells (Figure S3C) and NK cells (Figure S3D). Both of these PGD2-mediated inhibitory effects were reverted by OKN4395 treatment, to a similar extent that other DP1 inhibitors (Figure 3C-D). Overall, we were able to demonstrate using a series of distinct and complementary *in vitro* primary immune cell assays that OKN4395 alleviates both PGE2- and PGD2-induced suppression of classical immune cell functions through specific blockade of EP2, EP4 and DP1 receptors. We next seek to understand and support the rationale for combination with anti-PD1 therapies.

**Figure 3:**
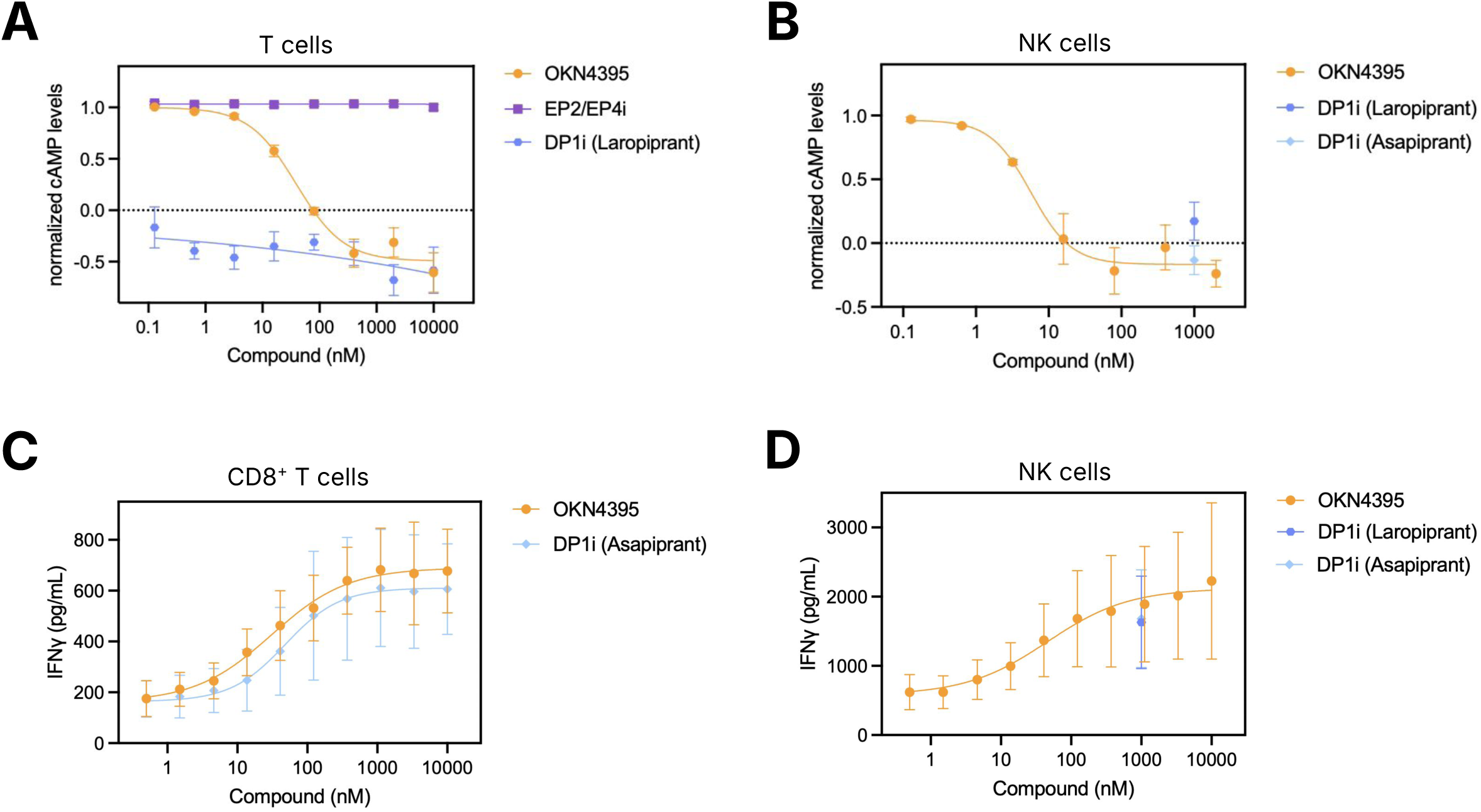
OKN4395 Alleviates PGD2- Induced Suppression of Immune Cell Functions In Vitro. (**A**) cAMP production of Pan T cells in the presence of PGD2 (EC90, 91.9nM). Values represent mean of n=3 healthy donors ± SEM. (**B**) cAMP production of NK cells in the presence of PGD2 (EC90, 15nM). Values represent mean of n=3 healthy donors ± SEM. (**C**) IFNγ secreted by CD8^+^ T cells stimulated with CD3/CD28 Dynabeads in the presence of PGD2 (200nM). Values represent mean of n=3 healthy donors ± SEM. (**D**) IFNγ secreted by NK cells stimulated with IL-2, IL-12, IL-15 and IL-18 in the presence of PGD2 (100nM). Values represent mean of n=3 healthy donors ± SEM.

### Combined OKN4395 and anti-PD1 blockade restores immune function and promotes antitumor activity

First, we wanted to understand whether OKN4395 alone or in combination with anti-PD1 could modulate IFNγ secretion from T cells cultured in a Mixed Leukocyte Reaction (MLR), an assay commonly used to assess PD-1/PD-L1 inhibitors. We established that in presence of allogeneic monocyte-derived dendritic cells (moDC), T cells were activated and that baseline IFNγ levels were detected in comparison to T cell cultured alone (Figure 4A). As previously published^34^, using pembrolizumab in order to block PD1, we were able to further induce IFNγ production. In these settings, addition of PGE2 was able to fully inhibit IFNγ production whether pembrolizumab was present or not. In these conditions, OKN4395 had a significant effect in reverting PGE2-mediated suppression of IFNγ. This effect was further amplified when OKN4395 was combined with pembrolizumab, reverting the suppression mediated by both PD1 and PGE2 communication channels. Interestingly, similar suppressive effects were observed when PGD2 was used instead of PGE2 confirming potent inhibitory functions of PGD2 (Figure 4B). As demonstrated for PGE2, OKN4395 enabled pembrolizumab induced IFNγ production in these PGD2-supplemented MLR settings.

**Figure 4:**
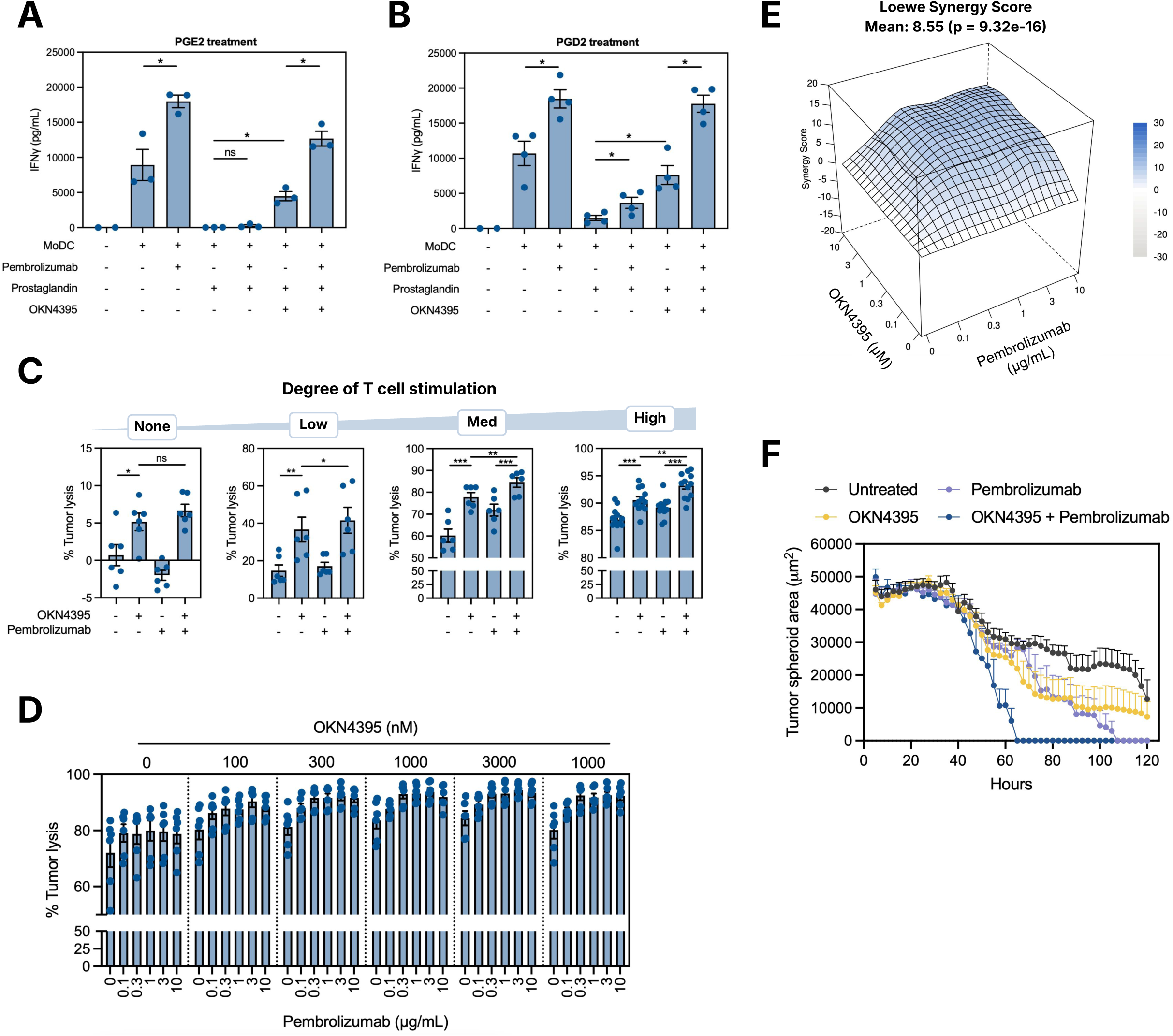
Combination of OKN4395 with anti-PD1 showed enhanced T cell activation and at least additive T cell-mediated tumor cell killing in vitro. (**A-B**) IFNγ secreted by Pan T cells co-cultured with MoDC in the presence of (A) PGE2 (500nM) or (B) PGD2 (1000nM). Datapoints represent individual IFNγ levels (PGE2: n=3, PGD2: n=4) with barplot representing the mean ± SEM. Significance was calculated using a paired t test. (**C**) T cell-induced tumor killing after 2 days of 2D co-culture. OKN4395 (10µM) and Pembrolizumab (10µg/mL) were added as single agents or in combination. Degree of T cell stimulation (None, Low, Med, High) corresponds to distinct levels of polyclonal activation. Values represent mean ± SEM of n=3 (None, Low, Med) and n=6 (High) healthy donors assessed in technical duplicates. Significance were calculated using a paired t test. (**D**) T cell-induced tumor killing after 2 days of 2D co-culture. OKN4395 and Pembrolizumab were added as single agents or in combination at the indicated concentrations. Values represent mean ± SEM of n=3 healthy donors assessed in technical duplicates. (**E**) Isobologram of OKN4395-Pembrolizumab additivity as assessed in 2D T cell-induced tumor killing. Loewe synergy score was calculated using the SynergyFinder+ online resource. (**F**) Tumor spheroid area over time assessed in a 3D T cell-induced tumor killing. OKN4395 (10µM) and Pembrolizumab (10µg/mL) were added as single agents or in combination. Values represent mean + SEM of n=3 healthy donors assessed in technical duplicates.

To further study potential interactions between anti-PD1 blockade and OKN4395 activity, we set up a killing assay co-culturing a bladder carcinoma cell line (5637 tumor cells) with primary total CD3^+^ T cells. 5637 cells were selected because of their natural expression of PGE2 (Figure S4A) and PD-L1 (Figure S4B). For this experiment to induce baseline killing of cancer cells a standard T-cell polyclonal activation was used. First, In absence of T-cell activation, as expected, no major killing of the 5637 cells was observed, even in presence of pembrolizumab or OKN4395 where it remained ≈5% in average (Figure 4C). Then using low, medium or high T-cell polyclonal activation, we observed a dose-dependent killing of 5637 cells mediated by T cells as a baseline, validating our assay. First, studying the low activation dose, we determined that only OKN4395 was able to increase tumor killing but that pembrolizumab had no effect. On the contrary, for higher doses of T cell activation, both pembrolizumab and OKN4395 had similar standalone activity and displayed a significantly increased killing activity once combined. In order to further quantify this potential interaction, we performed a double titration of both OKN4395 and pembrolizumab in this tumor killing assay (Figure 4D). Using the Loewe score as a metric to assess potential synergistic effects^35^, a significant positive score of 8.55 was calculated (Figure 4E). This was further confirmed using the HSA score as a distinct statistical methodology (Figure S4C). Finally, using similar experimental conditions than in the 2D tumor killing assay, a 3D assay was developed to follow upon time the effect of OKN4395 combined with pembrolizumab on the spheroid formed by 5637 tumor cells. This assay revealed that OKN4395 alone had similar effect than pembrolizumab in promoting killing of the tumor spheroid and that their combination could significantly accelerate the killing mediated by T cells of the tumor spheroid (Figure 4F, Movie S1).

### OKN4395 exhibits dose-dependent efficacy and synergistic potential with anti-PD1 *in vivo*

Next, we evaluated the *in vivo* pharmacology of OKN4395 as a single agent or in combination with pembrolizumab in the breast cancer EMT6 model. For this study, 1 million EMT6 cells were injected subcutaneously to the right flank of Balb/c mice. At day 7 after injection, OKN4395 b.i.d treatment was evaluated at three distinct doses, alone or in combination with anti-PD1 (Figure 5A). Single agent treatment revealed a significant tumor growth inhibition (TGI) proportional to the treatment dose. Indeed, OKN4395 at 35 mg/kg led to TGI=34% (p=0.21), while the 75 mg/kg and 150/125 mg/kg led to TGI=66% (p=0.004) and TGI=84% (p<0.0001), respectively (Figure 5B). Significantly, a body weight loss was observed at day 7 post treatment in the 150 mg/kg b.i.d single agent arm, which led to a dose reduction to 125 mg/kg for the second half of the treatment duration after 2 days of treatment holiday (Figure S5A). A similar effect of OKN4395 150mg/kg was observed on body weight when combined with anti-PD1, without additional impact of the combo on toxicity profile (Figure S5B). No significant body weight loss was observed in the 75 mg/kg or 35 mg/kg arms. Interestingly, within the 75 mg/kg and 150/125 mg/kg OKN4395 single arms, 2 Complete Response (CR) and 3 Partial Response (PR) could be observed illustrating the potential for OKN4395 to not only actively prevent tumor growth but also to lead to significant tumor reduction and shrinkage in some individuals (Figure 5C). In the EMT6 model, a twice weekly dose of anti-PD1 led to initial tumor reduction. Interestingly, the combination of anti-PD1 with OKN4395 led to potential increase in TGI compared to OKN4395 monotherapy (Figure 5B-D). Most importantly, combined treatment with anti-PD1 significantly increased the number of CR observed across the OKN4395 dose levels, with a total of 11 mice over 30 achieving CR (37%). These results are consistent with tumor volume measured at experiment termination (Figure 5D). On the last study day, blood of the experimental mice was collected to measure plasma drug concentration. In this *in vivo* efficacy study an approximate pharmacokinetic profile of the 3 dose levels of OKN4395 was determined at the last day drug administration. The OKN4395 plasma concentration between the two cohorts treated with the same dose (with and without anti-PD1) were comparable at any given time point, differing not more than by a factor of ∼2 (Figure S5C).

**Figure 5:**
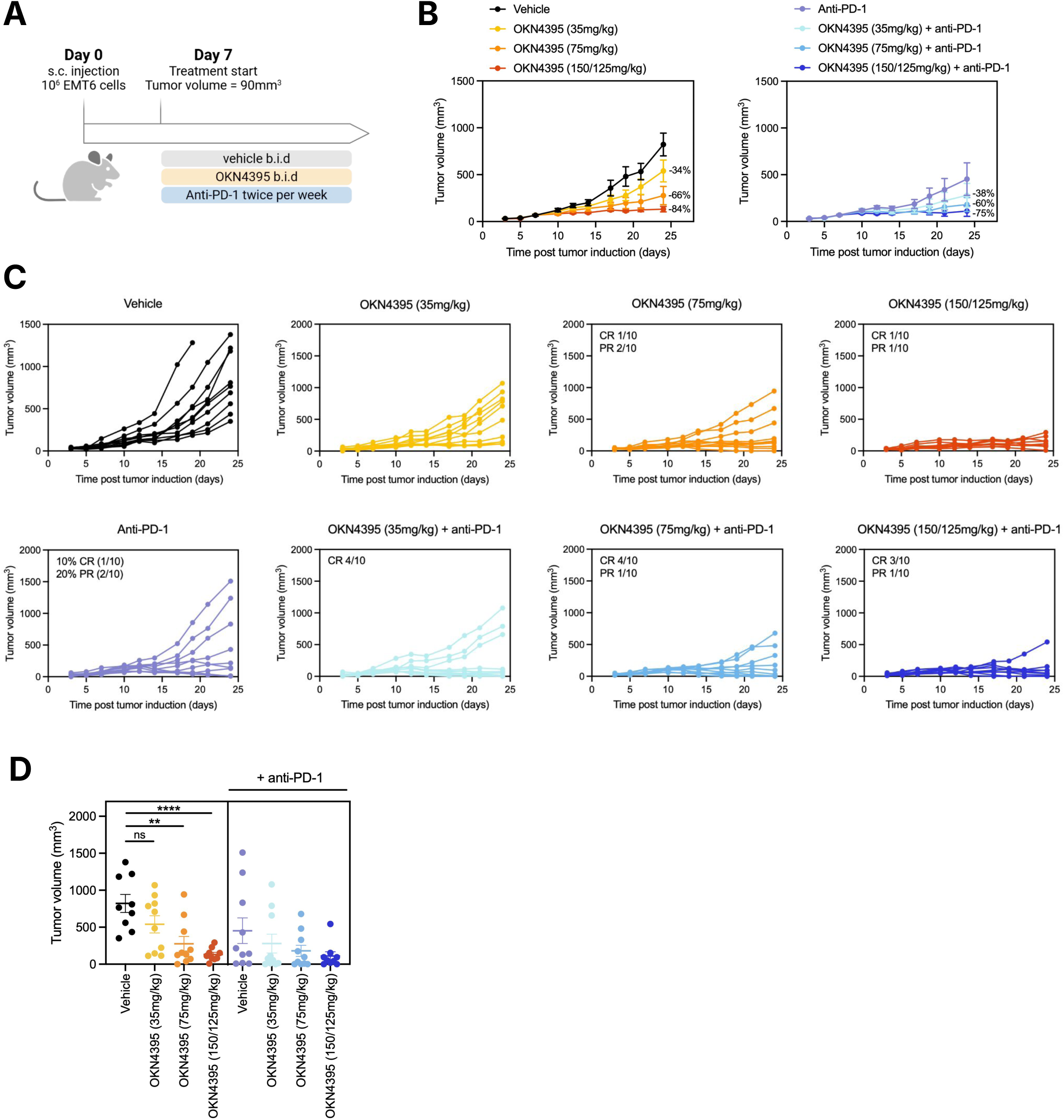
OKN4395 demonstrated high EMT6 tumor growth inhibition in vivo, alone or in combination with anti-PD1. (**A**) In vivo experimental scheme. Vehicle and OKN4395 were administered twice per day. Anti-PD1 was administered twice per week. (**B**) Tumor volume over time is shown for the indicated treatment. The left graph shows tumor control in response to treatment with vehicle or increasing OKN4395 concentrations. The right graph shows tumor control in response to treatment with anti-PD1 alone or in combination with increasing OKN4395 concentrations. The percentages indicate the relative difference in mean tumor volume on day 24 compared to the control group (left) or anti-PD1 group (right). (**C**) Individual animal tumor size graphs over time. Number of partial response (PR) and complete response (CR) were indicated for each treatment condition. (**D**) Individual tumor volume measured on day 24 (17 days after treatment initiation with the indicated treatments and doses). Values represent mean ± SEM of n=9-10 animals per group. A Mann-Whitney test was applied for statistical comparison of two groups.

## DISCUSSION

Overall our study demonstrates the unique pharmacologic profile of OKN4395 as an equipotent EP2, EP4 and DP1 antagonist with high specificity and potent *in vitro* and *in vivo* antitumoral functions. To our knowledge, OKN4395 is a first-in-class therapeutic with the potential to address several challenges associated with the therapeutic modulation of COX2 pathway in solid tumors.

Strategies for blocking PGE2 production by targeting the upstream COX enzymes, while promising, have demonstrated toxicities that have limited clinical use as anti-cancer agents^20^. A recent mitigating approach aims at blocking only those specific downstream PGE2 receptors involved in suppressing antitumor immunological responses e.g. EP4, to spare other pathways essential for normal tissue homeostasis^21,22^. Recently, some of these more targeted strategies have revealed interesting signs of clinical efficacy^25^. There is, however, mechanistic evidence demonstrating that most of the antitumoral activity mediated by EP4 blockade could be compensated by PGE2 mediated immunosuppressive signaling through EP2^26,27^ with the result that EP4/EP2 dual inhibitors have also been investigated clinically^26,36^. OKN4395 is a balanced, equipotent EP2 and EP4 antagonist, a mechanism supporting robust blockade of PGE2-mediated signaling. Further, OKN4395 not only equipotently blocks EP2 and EP4 but also brings equipotent blockade of DP1, a well known receptor of the PGD2 ligand with mechanistically convergent downstream second messenger signaling.

The role of DP1 in oncology is less established than that for EP2 and EP4. However previous studies demonstrated that cAMP was the main second messenger induced by the binding of PGD2 to DP1, which is mechanistically well-aligned with the downstream effector mechanism of PGE2 signaling through EP2 and EP4^37^. The parallel approach of blocking cAMP production caused by PGE2 signaling through EP2/EP4 and also through PGD2 signaling through DP1 is important because cAMP mediated signaling, specifically in immune cells, has been recognized as a central mechanism of immune suppression^38^. In addition, preliminary *in vitro* studies investigating the role of PGD2 on immune cells suggested that DP1 agonism could lead to decreased IFNγ secretion in either T or NK cells, in a similar fashion than what is observed with PGE2^30,39^. The potential for PGE2 blockade to be bypassed by PGD2-mediated DP1 signaling provides a rationale for triple inhibition of EP2, EP4 and DP1.

The role of PGD2 in tumors is coming under increasing mechanistic investigation. For example, HPGDS, a main synthase of PGD2, has been recently shown to be significantly associated with macrophage infiltrate and with resistance to anti-PD1 in a melanoma human clinical study^29^. The authors further demonstrated the potential pro-tumoral role of the PDG2-DP1 axis in elegant *in vitro* and *in vivo* models with proof-of-concept data using genetic and pharmacologic ablation of DP1 expression.

The effect of OKN4395 combined with PD-1 inhibition opens an intriguing path forward in next-generation immunomodulatory cancer therapies. Importantly, several initially promising novel immuno-oncology therapies when combined with anti-PD(L)1 checkpoint inhibition have not provided the hoped-for step change in clinical outcomes^40,41^. Many of these novel agents exclusively targeted specific mechanisms on T cells, missing potentially important antitumor contributions from other cell types. In contrast, by interrupting the COX2-PGE2 axis OKN4395 modulates a broad range of innate and adaptive immune cells—shaping a pro-inflammatory, antitumoral state in NK, CD8^+^ and CD4^+^ T cells—and also influences dendritic cells, neutrophils, and macrophages in the myeloid compartment. Engagement of the broader immune system in this manner with OKN4395 combined with checkpoint inhibitors is a promising additional avenue for clinical exploration.

## METHODS

### Human PBMCs isolation

Apheresis blood samples from healthy human donors were obtained from the Etablissement Français du Sang (EFS) after informed consent, in conformity with EFS ethical guidelines. Peripheral blood mononuclear cells (PBMCs) were isolated by centrifugation on a density gradient (Lymphoprep layered into Sepmate tubes, StemCell Technologies).

### cAMP assay for primary human immune cells

T cells and NK cells were isolated from frozen healthy donor PBMCs by magnetic isolation with the Pan T cell and NK cell isolation kit (130-096-535 and 130-092-657 respectively, Miltenyi Biotech). Cells were plated at 2×10^5 cells/well and incubated with inhibitors, as indicated in the results, for 20min at 37°C. Subsequently, PGE2 (2296/10, Biotechne) or PGD2 (12010, Cayman) were added and cells were incubated for 30min at 37°C. cAMP levels were measured using the cAMP-Glo™ Assay (V1502, Promega) following manufacturer’s instructions and luminescence levels were measured with the GloMax® Discover Microplate Reader (GM3000, Promega). Normalized cAMP levels = (x - negative control) / (positive control - negative control); negative control is DMSO, positive control is highest concentration of PGE2 or PGD2, respectively.

### cAMP assay for cell lines naturally and exogenously expressing EP2, EP4 or DP1 receptors

Human glioblastoma SF295 and human breast BT549 cancer cell lines, which endogenously express either EP2 receptor or EP4 receptor, respectively, were used. Stable HEK293 cell clones overexpressing either human EP2 (hEP2, HGNC: 9594) or EP4 (hEP4, HGNC: 9596) or IP (hIP, HGNC: 9602) receptors or mouse EP2 (mEP2) or EP4 (mEP4) receptors have been established by transfection of HEK293 cell line (DSMZ, ACC305) with the following plasmids respectively: phCMV1.3HA.hEP2, phCMV1.3HA.hEP4, pcDNA3.hIP, phCMV1.3HA.mEP2 or phCMV1.3HA.mEP4 plasmid. HEK293 cell line was transduced by the respective DNA sequences carried by lentiviral vectors obtained from Vectalys for exogenous expression of either dog (canis lupus) EP2 (dEP2) or EP4 (dEP4) receptors or monkey (macaca fascicularis) EP2 (mkEP2) or EP4 (mkEP4) receptors or rat (rattus norvegicus) EP2 (rEP2) or EP4 (rEP4) receptors. Overexpressing human DP1 (hDP1, HGNC: 9591) CHO-K1 stable clones were established and selected following transfection of CHO-K1 cells with the pcDNA5/FRT.hDP1R plasmid. HEK293 (CRL-3216, ATCC) cells transiently expressing mouse DP1 (mDP1), rat DP1 (rDP1) or monkey DP1 (mkDP1) were generated by lipofectamine (L3000008, Thermo Fisher Scientific) transfection with the following plasmids respectively: pRP-EF1A.mDP1:P2A:EGFP, pRP-EF1A.rDP1:P2A:EGFP or pRP-EF1A.mkDP1:P2A:EGFP (Vector Builder). For HEK293 cells transiently expressing DP1 receptors, cells were harvested 48h after transfection and cAMP levels were measured as for primary human immune cells (see section above). As a detection method for cAMP on cell lines stably expressing the receptors, the HTRF cAMP Gs Dynamic Detection Kit (62AM4PEJ, Cisbio Cat/Revvity) was used as previously described^31,33^. EC80 of PGE2 (14010, Cayman) or PGD2 (PG-005, Enzo) or iloprost (18215, Cayman) were used as agonists as indicated in the figures. Reading was performed using a BMG LABTECH PHERA star reader (Excitation: 337 nm, Emission: 620 and 665 nm). HTRF ratio is calculated as follows: HTRF ratio = 10′000 ×(emission at 665 nm / emission at 620 nm). When indicated, data are expressed as “cAMP levels in percent control” calculated as follows: cAMP levels (% control) = (ratio (DMSOcontrol) − ratio (sample)) / (ratio (DMSOcontrol) – ratio (ligand at EC80)) × 100. IC50 values and curves were generated with GraphPad Prism dose-response four-parameter fit.

### PathHunter GPCR assays for cell lines exogenously expressing EP1 or EP3 receptors

Stable U2OS cell clones overexpressing human EP1 (hEP1, HGNC: 9593) and stable CHO-K1 cell clones overexpressing human EP3 (hEP3, HGNC:9595) were used. As a detection method for chemiluminescent detection of activated G-protein coupled receptors (GPCRs), assay PathHunter® Glo Detection kit (93-0001, DiscoverX) was used as described in the “large GPCR activity screen” section. EC80 of PGE2 (14010, Cayman) was used as an agonist as indicated in the figures.

### GPCR selectivity screen

The gpcrMAX and orphanMAX panels from Eurofins were used to identify potential interactions with a curated selection of known G-protein coupled receptors (GPCRs) and orphan GPCR targets. In these panels, OKN4395 was evaluated at two distinct concentrations (1µM and 10µM). Briefly, both panels are based on The PathHunter® β-Arrestin assay initially developed by DiscoverX monitoring the activation of a given GPCR in a homogenous, non-imaging assay format using a technology developed by called Enzyme Fragment Complementation (EFC) with β-galactosidase (β-Gal) as the functional reporter measured using chemiluminescent PathHunter® Detection Reagents. The full list of GPCR tested for OKN4395 activity can be found in Figure S1E. For antagonist-mode assays, cells were co-treated with the receptor’s natural ligand and OKN4395. Activity was quantified as the percentage of the natural ligand response in agonist-mode, and the percentage of reduced activity in antagonist-mode.

### OKN4395 and other small molecule inhibitors

OKN4395, described in WO2017085198A1, was synthesized as described therein. For experiment controls, the following inhibitors were used across several experiments: PF-04418948 (S7211, Selleckchem) is a selective EP2 receptor antagonist^42^. Palupiprant/E7046 (HY-103088, MedChemExpress) is a selective EP4 receptor antagonist^43^. Asapiprant (30008, Cayman) is a selective DP1 receptor antagonist^44^. Laropiprant (10009835, Cayman) is a selective DP1 receptor antagonist^45^.

### T cell proliferation assay

Polyclonal CD3 T cells were isolated from frozen healthy donor PBMCs by magnetic isolation with the pan T cell isolation kit (130-096-535, Miltenyi Biotech). Cells were stained with 5µM of CellTrace CFSE (C34554, Thermo Fisher Scientific) following the manufacturer’s recommendation. Cells were seeded in 96 well plates at 5×10^4 cells/well. PGE2 (500nM) and respective inhibitors were added and T cells were stimulated with anti-CD3/CD28 Dynabeads (11132D, Thermo Fisher Scientific) at a bead:cell ratio of 1:10. After 5 days, CFSE dilution in CD4^+^ and CD8^+^ T cells was analyzed using anti-human CD8a-BV650 (301042, Biolegend) and anti-human CD4-BV605 (562658, BD Biosciences) antibodies and MACSQuant analyzer 16 flow cytometer. The percentage of proliferating T cells was calculated as 100 x (sample - PGE2 treatment)/(no PGE2 treatment - PGE2 treatment).

### Flow cytometry

PD-L1 expression on 5637 tumor cells was analyzed by flow cytometry following staining with BV650 anti-human PD-L1 (clone 29E.2A3, Biolegend) antibody. In all flow cytometry experiments, cell viability was assessed using fixable viability dye eFluor 780 (65-0865-14, Thermo Fisher). Samples were acquired on a MACSQuant Analyzer 16 (Miltenyi). Data was analyzed using FlowJo software (Becton, Dickinson & Company).

### Quantification of PGE2 by ELISA

5637 tumor cells were plated at 0.125×10^6 cells/mL in a 6-well plate and incubated for 72 hours at 37°C. PGE2 concentration in the supernatant was determined by ELISA (KGE004B, Biotechne) following manufacturer’s recommendations.

### Tumor killing assay

5637 tumor cells were transduced (MOI of 10) with lentiviral particles encoding for eGFP, Firefly Luciferase and Puromycin resistance. Transduced cells were selected using 2µg/mL Puromycin (Merck, P4512). Polyclonal CD3 T cells were isolated from frozen healthy donor PBMCs by magnetic isolation with the pan T cell isolation kit (130-096-535, Miltenyi Biotech). For 2D tumor killing assays, 5637 tumor cells were seeded at 5×10^3 cells/well in 96 well plates. After 24 hours, T cells were added at 5:1 effector:target ratio. After 2 days, tumor viability was assessed by addition of 0.3mg/mL per well of D-Luciferin (122799, Revvity) and luminescence measurement using GloMax Discover System (Promega). The percentage of specific tumor lysis was calculated as 100 - (% live tumor cells) where % live tumor cells corresponds to 100 x (sample - background)/(negative control - background).

For 3D tumor killing assays, 5637 tumor cells were seeded at 1×10^4 cells/well in 96 well Ultra-Low Attachment (ULA) plates. After 24 hours, T cells were added at 5:1 effector:target ratio in the presence of 10µM OKN4395 and 10µg/mL Pembrolizumab (aRMP Keytruda, Evidentic). Plates were incubated inside a CellVoyager CQ1 high-content analysis system (Yokogawa), during which live images were acquired every 2.5 hours. Tumor spheroid area was assessed over time using a maximum intensity projection.

### *Ex vivo* whole blood assay

Heparinized blood from healthy donors was added to TruCulture tubes (Rules Based Medicines) previously spiked with LPS or LPS + PGE2 (14010, Cayman) at different concentrations and the indicated concentrations of OKN4395. Tubes were incubated in dry block thermostats at 37°C. After 24 hours, supernatants were harvested, aliquoted and stored at −20°C. Cytokine concentrations were assessed via Luminex-based Multi-Analyte Profiles to evaluate immune cell activities.

### Functional readout with CD8^+^ T cells and NK cells

CD8+ T cells or NK cells were isolated from frozen healthy donor PBMCs by magnetic isolation with the CD8 T cells or NK cell isolation kit (130-096-495 and 130-092-657, Miltenyi Biotech). Cells were plated in 96 well plates at 1×10^5 cells/well and incubated with the respective inhibitors (EP2 inhibitor: PF-04418948, S7211, Selleckchem; EP4 inhibitor: Palupiprant/E7046, HY-103088, MedChemExpress; DP1 inhibitors: Asapiprant, 30008, Cayman and Laropiprant/MK-0524, 10009835, Cayman; OKN4395) for 20 min. PGE2 or PGD2 were added and CD8+ T cells were incubated for 30min before stimulation with anti-CD3/CD28 Dynabeads (11132D, Thermo Fisher Scientific) at a bead:cell ratio of 1:1 for 8h, NK cells were directly stimulated with IL-2, IL-12, IL-15 (130-097-745, 130-096-705, 130-095-764, Miltenyi Biotech) and IL-18 (9124-IL-050/CF, biotechne), each at a concentration of 10ng/ml for 6h. IFNγ levels in the cell culture supernatants were measured by U-PLEX assay (K15067M-2, MSD) or Lumit Immunoassay (W6041, Promega).

### Co-culture of MoDC and T cells

Monocytes were isolated from fresh healthy donor PBMCs using CD14 microbeads (130-050-201, Miltenyi Biotech) and cultured for 6-7 days with 50ng/ml IL-4 (130-094-117, Miltenyi Biotech) and 10ng/ml GM-CSF (130-093-867, Miltenyi Biotech). Differentiated MoDCs were cultured at a 1:10 MoDC:T cell ratio with allogeneic Pan T cells isolated from frozen healthy donor PBMCs in the presence of 5ug/ml anti-CD3 antibody (16-0037-85, Thermo Fisher Scientific), 500nM PGE2 (2296/10, biotechne) / 1000nM PGD2 (12010, Cayman) / 10µM OKN4395 / 10 µg/ml Pembrolizumab (aRMP Keytruda, Evidentic) for 6-7 days. IFNγ levels in the cell culture supernatants were measured by Lumit assay (W6041, Promega).

### RT-qPCR

RNA was extracted from CD4^+^ T cells, CD8^+^ T cells or NK cells that had been isolated from healthy donor PBMCs or from HEK293T cells (CRL-3216, ATCC) using the RNeasy Mini kit (74106, Qiagen). RNA was reverse transcribed using the High-Capacity cDNA Reverse Transcription kit (4374967, Fisher Scientific). qPCR was performed in a QuantStudio 7 Pro using the TaqMan Fast Advanced Master Mix (4444557, Fisher Scientific) and the following TaqMan assay ID (Thermo Fisher Scientific) specific for the respective human genes of interest: PTGER1 (Hs00168752_m1), PTGER2 (Hs00168754_m1), PTGER3 (Hs00168755_m1), PTGER4 (Hs00168761_m1), PTGDR (Hs00235003_m1), PTGDR2 (Hs00173717_m1). Relative gene expression was calculated as 2^-(Ct(gene of interest)-Ct(GAPDH)).

### Docking model

#### Structure analysis and preparation

7CX3, an agonist-bound structure was selected to represent EP2; 6AK3 was selected to represent EP3; 5YWY, an antagonist-bound structure was selected to represent EP4. DP1 structure was downloaded from AlphaFoldDB^46^. Structures were prepared using Schrödinger’s Maestro v14.2 using a pH of 7.4. The prepared structures were minimized to maximum 0.3 RMSD from the input and aligned. The DP1 structure being an apo structure, the binding site presents a close conformation, requiring the use of induced-fit docking to successfully identify binding poses for our compound series.

#### Docking

The compounds to dock were preprocessed using the LigPrep protocol (Schrödinger’s Maestro v14.2). For each protein, the docking grid was defined on the binding pocket defined after aligning all receptors. Constraints were added to encourage salt bridge formation with the pocket’s arginine residue (Arg302 in EP2, Arg333 in EP3, Arg316 in EP4 and Arg310 in DP1). Prepared ligands were docked in the prepared grids using Glide. For DP1, due to the closed conformation of the structure, an Induced-Fit Docking protocol was used with OKN4395 only. The resulting best pose was chosen to define the DP1 receptor, with a new docking grid centered on OKN4395, and to dock all other molecules with Glide SP. Output docking poses were manually checked for all ligands, prioritizing the presence of the salt bridge between the carboxylic acid and the arginine residue in the pocket.

#### Molecular Dynamics

to further explore the stability and possible rearrangement of residues, we ran molecular dynamics simulations on EP3, EP4 and DP1-based complexes. The complexes were put in a bilayer membrane by aligning the complexes on the Z axis and center them on the XY plane. Then, the resulting coordinate files were used with openMM^47(p8)^ and openFF^48^ to construct the relevant system: a POPC bilayer, a water and ions box (NaCl at 130mM) and the protein-ligand complex. To equilibrate the system, the backbone of the protein was frozen and the system warmed from 0 to 310 K in a NVT simulation, by 10 K increments. Following the heating, a short NPT equilibration using a Monte Carlo anisotropic barostat (1 bar of pressure, surface tension of 200 bar/nm) was done while gradually lifting the restraints on the backbone. Frames were written every 0.1 ns for the 3 runs of 50 ns of NPT production (3 x 50 ns).

### *In vivo* mouse experiments

Seven weeks old female Balb/cByJ mice were obtained from Charles River. The *in vivo* study was conducted at Oncodesign (Dijon, France) whose animal care unit is approved by French ministries of Agriculture and Research (agreement number A21231011EA). On day 0, tumors were established by subcutaneous injection of 1×10^6 EMT6 cells (CRL-2755, ATCC) in the right flank of the mice. On day 7, therapeutic treatment started after animal randomization. OKN4395 was formulated in an aqueous solution of 50mM citrate buffer, pH 4.0 containing 0.5% methylcellulose and administered p.o. twice daily at 35, 75 or 150mg/kg initially. After seven days of consecutive treatment with 150mg/kg, the mice were allowed to recover on days 15 and 16 before treatment was resumed with 125mg/kg twice daily. Twice daily oral administration of the vehicle alone served as control. Anti-PD1 antibody (BE0146, clone RMP1-14, BioXcell) was diluted in PBS to a final concentration of 1mg/mL and injected i.p. at a volume of 10mL/kg body weight. Injections with PBS alone served as control. Administration started on day 7 and occurred twice per week for two consecutive weeks. Mouse behavior was recorded every day. Body weights were measured twice a week. The length (L) and width (W) of tumors were measured twice a week with calipers and tumor volume (V) was calculated by the formula V = (W x W x L) / 2.

### Plasma bioanalysis of OKN3595 in mice

Plasma levels of OKN4395 in EMT6 tumor-bearing mice were analyzed on day 28 after twice-daily administration of 35, 75 and 150/125 mg/kg. Plasma concentrations were determined using protein precipitation and LC-MS/MS (liquid chromatography coupled to tandem mass spectrometry). The bioanalytical method was fully validated and used a deuterated OKN4395 as internal standard.

### Quantification and statistical analysis

All statistical tests were performed using Prism v.10.6.0 (GraphPad). Data are expressed as mean ± SEM as indicated in figure legends. Paired t-tests and Mann-Whitney U tests were used as indicated in individual figure legends. All statistical tests were two-tailed with a significance level of 0.05. ns, not significant; *p<0.05; **p<0.01; ***p<0.001; ****p<0.0001.

## ACKNOWLEDGEMENTS

The authors wish to deeply thank Benno Schindelholz, Bénédicte Haenig, Marcel Keller from Idorsia Pharmaceutical for the help provided in sourcing and transferring data and material that were essential for this study. The authors also wish to thank Dominique Meyer, Michael Török and Enrico Vezzali from Idorsia Pharmaceutical for their valuable contributions to *in vivo* studies. The authors thank Anna Song for technical help provided in computational modeling. Generative artificial intelligence was used to revise and improve the clarity of text written by the authors.

---

**Figure 1S:**
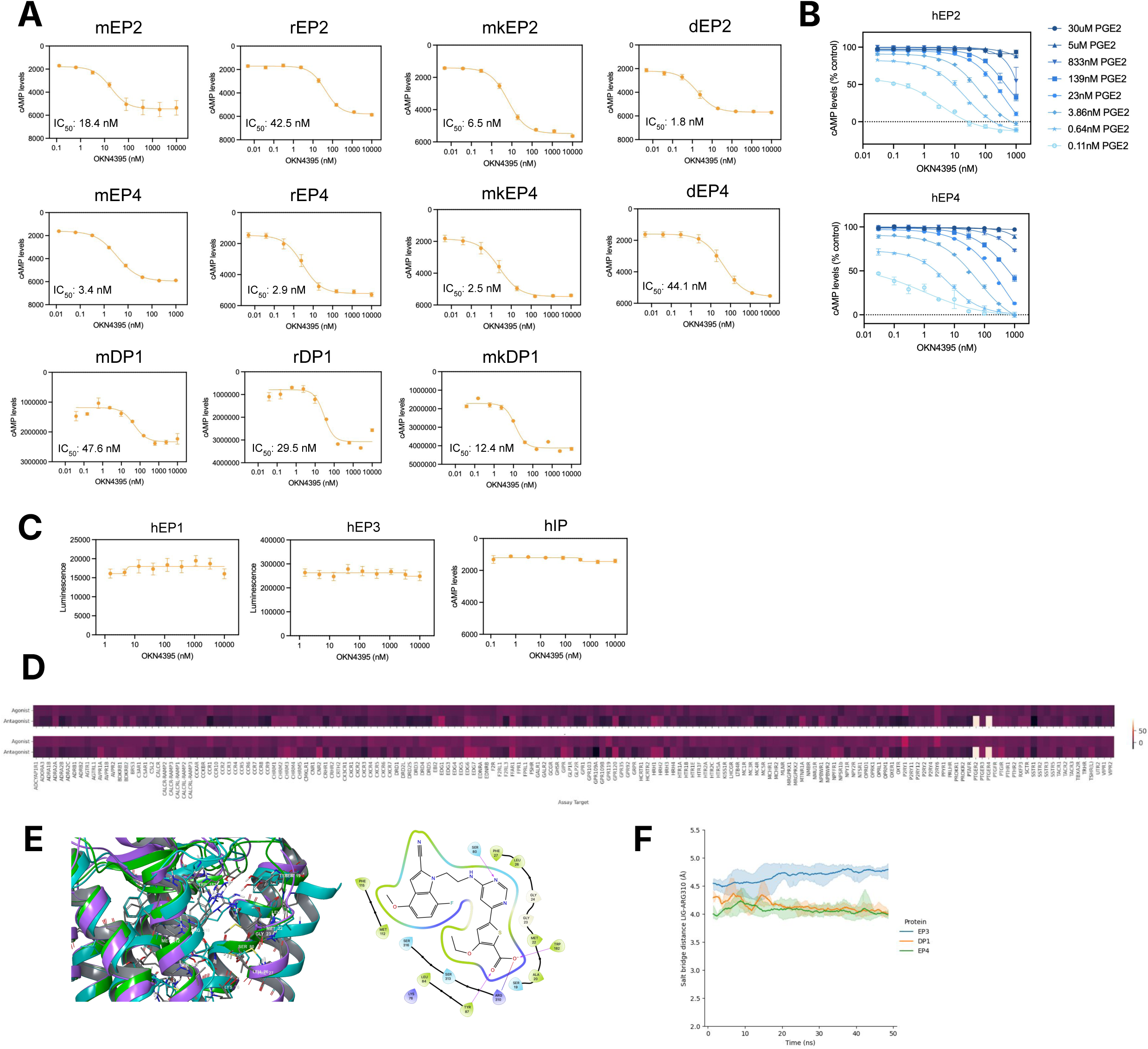
Supplementary data associated to figure 1. (**A**) cAMP production after stimulation with PGE2 (EC80) measured on HEK293 stable clones expressing EP2 or EP4 from different species or stimulation with PGD2 (EC80) measured on transfected HEK293 cells transiently expressing DP1 from different species. EC80 PGE2 concentrations are mouse EP2 (mEP2) 300pM, mouse EP4 (mEP4) 200pM, rat EP2 (rEP2) 35-45pM, rat EP4 (rEP4) 5.8-7.8pM, monkey EP2 (mkEP2) 35-40pM, monkey EP4 (mkEP4) 4.2-7.8pM, dog EP2 (dEP2) 65-80pM and dog EP4 (dEP4) 17.8-22.7pM. EC80 PGD2 concentrations are mouse DP1 (mDP1) 2.5nM, rat DP1 (rDP1) 4.4nM and monkey DP1 (mkDP1) 0.75nM. Values represent mean of n=4 (mEP2), n=6 (mEP4), n=10 (rEP2, rEP4, mkEP2, mkEP4, dEP2, dEP4), n=2 (mDP1, rDP1, mkDP1) technical replicates ± SEM. (**B**) cAMP production after stimulation with different PGE2 concentrations measured on SF295 cells (left) or BT549 cells (right) naturally expressing EP2 and EP4 receptor respectively. Values represent mean of n=2 technical replicates ± SEM. (**C**) Activation of (left) hEP1 and (middle) hEP3 after stimulation with PGE2 (EC80, 45nM and 15nM respectively) was measurement on U2OS cells and CHO-K1 cells expressing human EP1 and human EP3 respectively. (Right) cAMP production after stimulation with iloprost (EC80, 60pM) measured on HEK293 stable clones expressing hIP. Values represent mean of n=8 (hEP1, hEP3) and n=4 (hIP) technical replicates ± SEM. (**D**) Selectivity of OKN4395 (top 1µM; bottom 10µM) across 166 G protein-coupled receptors. OKN4395 activity was quantified as the percentage of the natural ligand response, with reduced activity indicating inhibition. (**E**) Superimposition of EP4 (cyan), EP2 (purple) and DP1 (green) binding sites (left). Interactions made by a representative of OKN4395 series in DP1 (right). All the main residues essential for the interactions are conserved in the three proteins. (**F**) Evolution of the distance between the carboxylate of OKN4395 and the interacting arginine residue in EP3, EP4 and DP1, showing the gradual weakening of its strength through movement of the ligand in EP3 and the stability in EP4 and DP1 (solid line: averaged across the triplicates, shaded area: standard deviation).

**Figure 2S:**
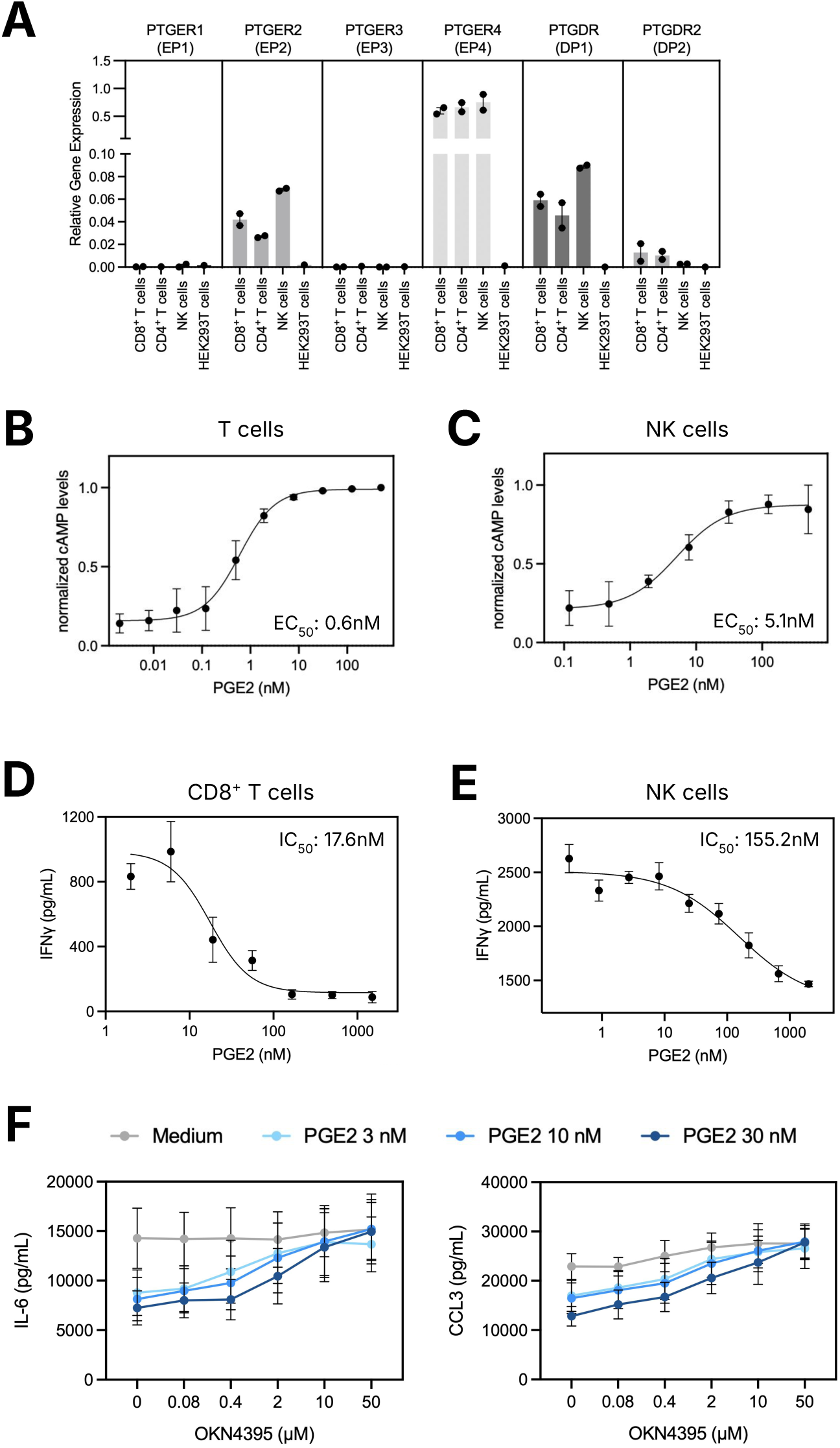
Supplementary data associated to figure 2. (**A**) Relative gene expression of prostaglandin receptors on blood derived primary cells or HEK293T cells. Values represent mean of n=2 healthy donors ± SEM and n=1 for HEK293T cells. (**B**) cAMP production of Pan T cells. Values represent mean of n=3 healthy donors ± SEM. (**C**) cAMP production of NK cells. Values represent mean of n=3 healthy donors ± SEM. (**D**) IFNγ secreted by CD8^+^ T cells stimulated with CD3/CD28 Dynabeads. Values represent mean of n=4 healthy donors ± SEM. (**E**) IFNγ secreted by NK cells stimulated with IL-2, IL-12, IL-15 and IL-18. Values represent mean of n=3 healthy donors ± SEM. (**F**) IL-6 and CCL-3 concentrations recovered in the supernatant of a human whole blood test engagement in the presence of LPS (0.1µg/mL) and increasing concentrations of PGE2 (0-3-10-30nM) for 24 hours. Values represent mean of n=3 healthy donors ± SEM.

**Figure 3S:**
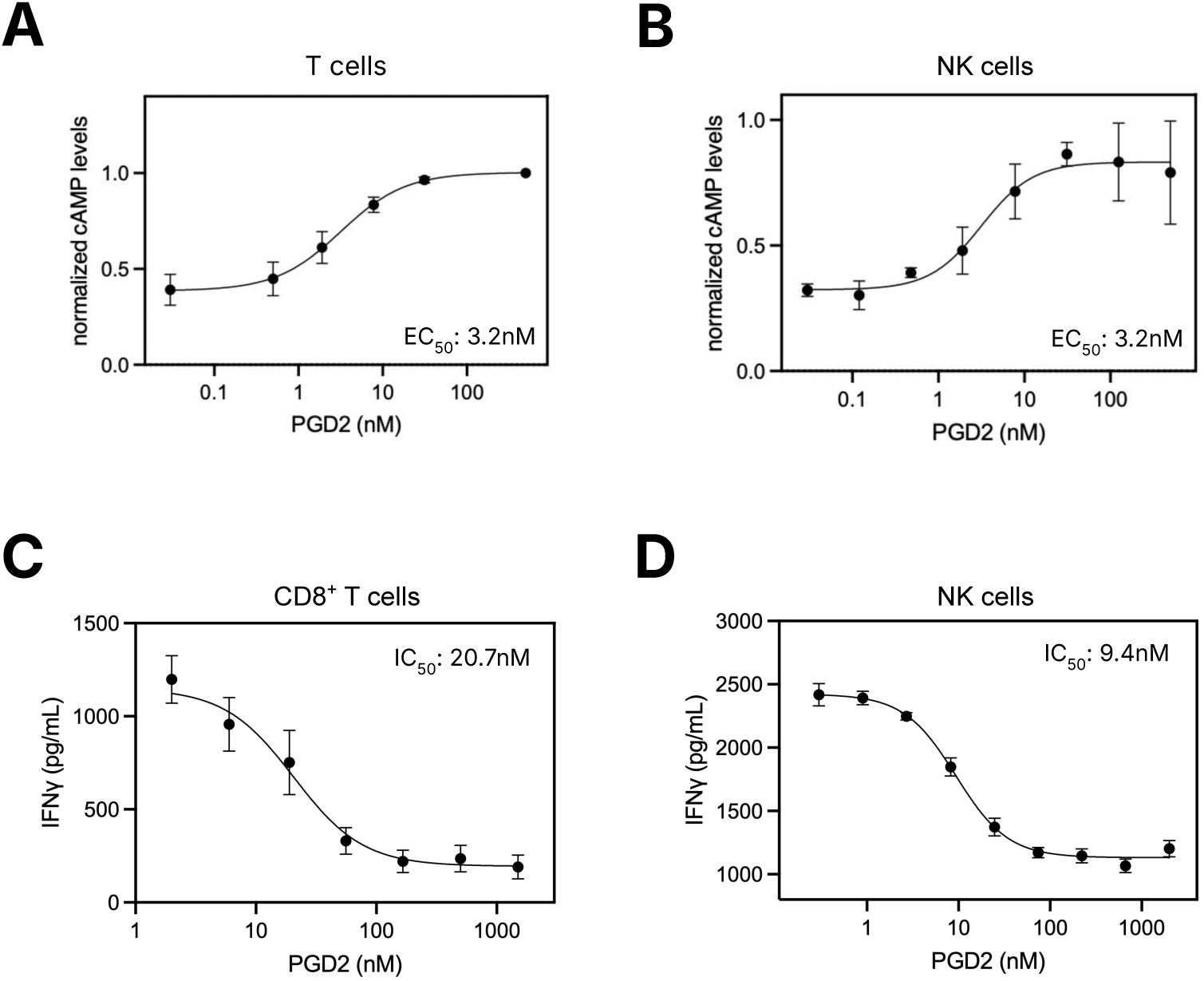
Supplementary data associated to figure 3. (**A**) cAMP production of Pan T cells. Values represent mean of n=3 healthy donors ± SEM. (**B**) cAMP production of NK cells. Values represent mean of n=3 healthy donors ± SEM. (**C**) IFNγ secreted by CD8^+^ T cells stimulated with CD3/CD28 Dynabeads. Values represent mean of n=4 healthy donors ± SEM. (**D**) IFNγ secreted by NK cells stimulated with IL-2, IL-12, IL-15 and IL-18. Values represent mean of n=3 healthy donors ± SEM.

**Figure 4S:**
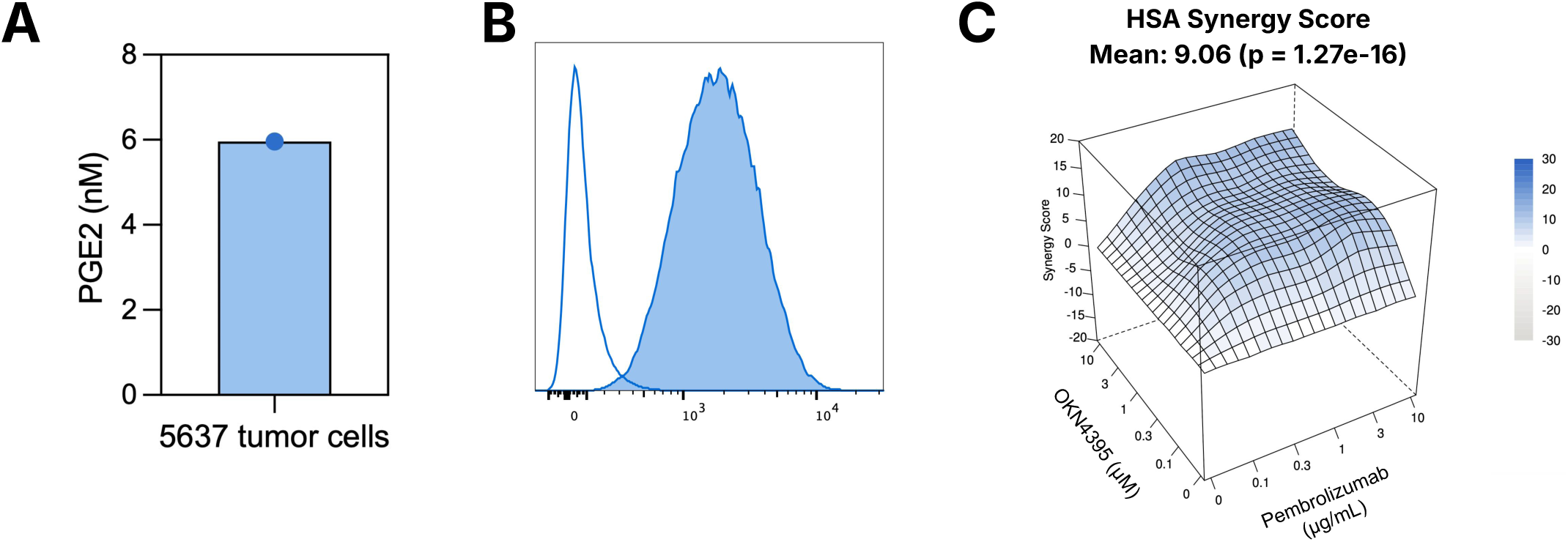
Supplementary data associated to figure 4. (**A**) Quantification of PGE2 concentration in the supernatant of 5637 tumor cells by ELISA. (**B**) PD-L1 surface expression by 5637 tumor cells assessed by flow-cytometry. Empty histogram corresponds to unstained 5637 tumor cells. (**C**) Isobologram of OKN4395-Pembrolizumab additivity as assessed in 2D T cell-induced tumor killing. HSA synergy score was calculated using the SynergyFinder+ online resource.

**Supplementary movie 1:**
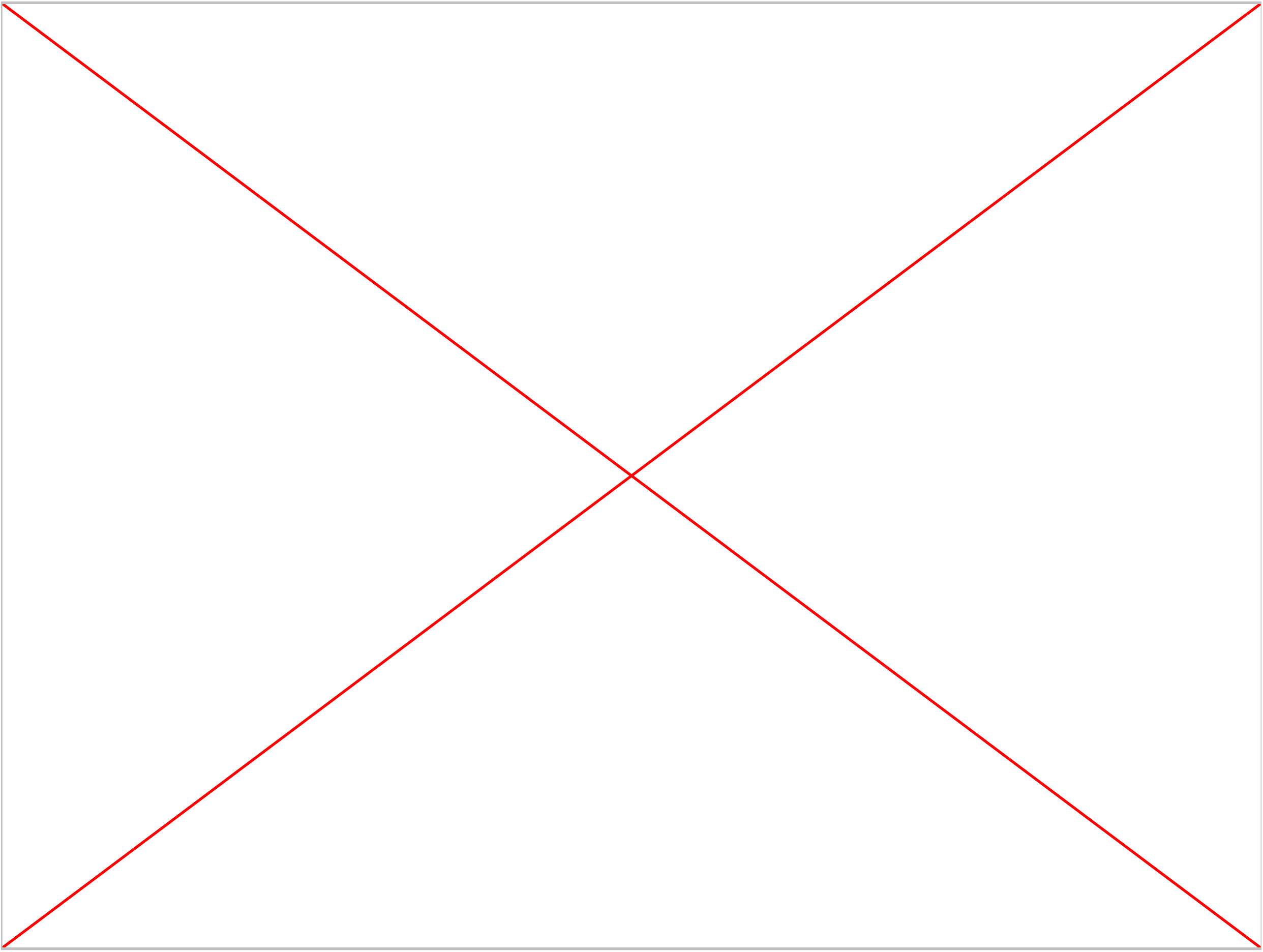
Combination of OKN4395 with anti-PD1 improves T cell efficacy in a 3D tumor killing assay. 5637 tumor cells expressing EGFP were co-cultured with T cells for 5 days. OKN4395 (10µM) and Pembrolizumab (10µg/mL) were added as single agents or in combination. Green area corresponds to tumor spheroid area assessed over time. Scale bars represent 350µm.

**Figure 5S:**
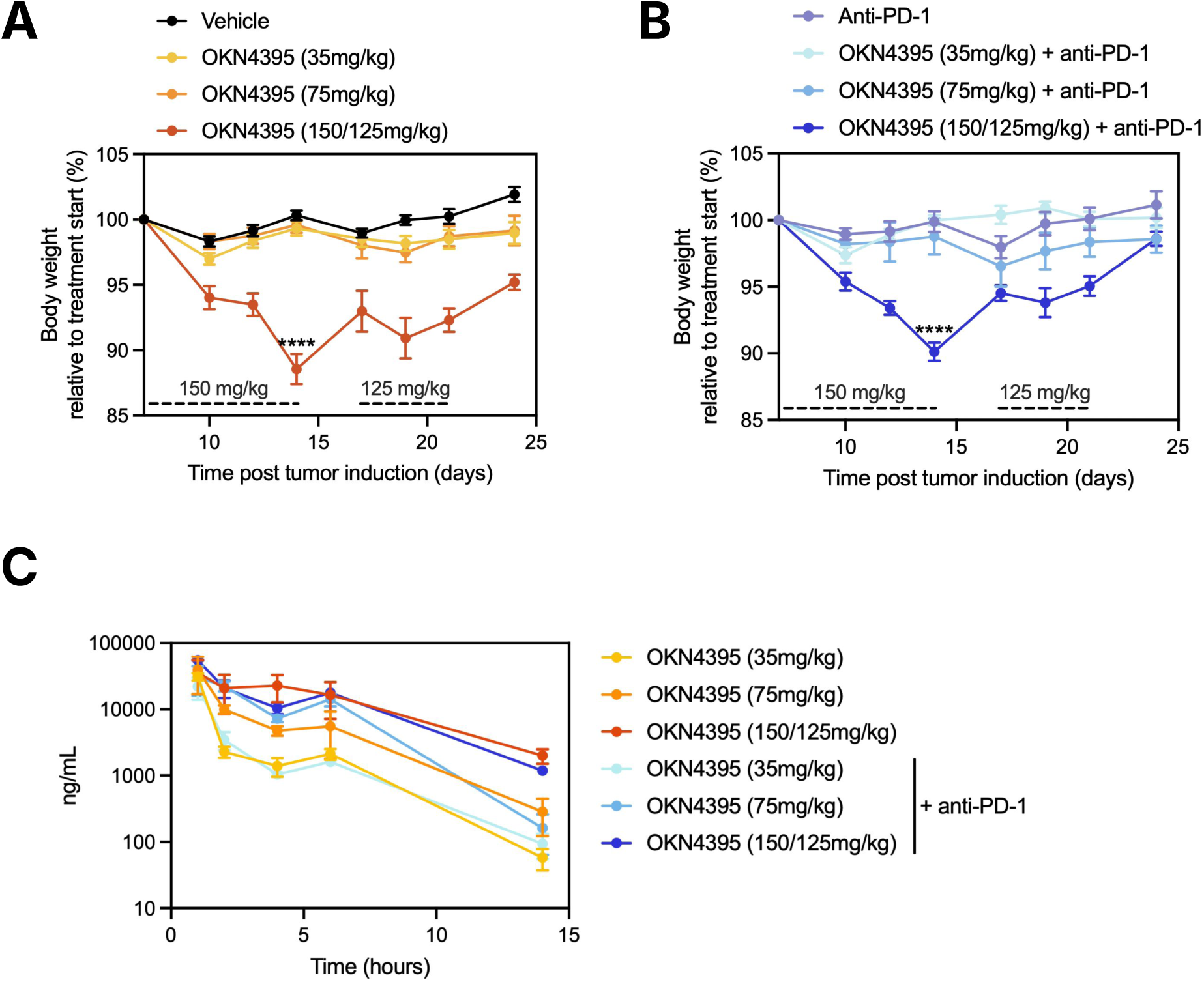
Supplementary data associated to figure 5. (**A-B**) Evolution of body weight relative to treatment start on day 7 is shown as percentage for animals treated with (A) vehicle or increasing OKN4395 concentrations and (B) anti-PD1 alone or in combination with increasing OKN4395 concentrations. Values represent mean ± SEM. The dashed lines mark the time frames of treatment in the high dose cohort with either 150 or 125 mg/kg. A Mann-Whitney test was applied for statistical comparison of two groups (vehicle *vs* OKN4395 150/125 mg/kg and anti-PD1 versus anti-PD1 + OKN4395 150/125 mg/kg on day 14). (**C**) OKN4395 plasma concentration on the final day of the study (day 28). Plasma of mice from the different treatment groups was collected 1h, 2h, 4h, 6h and 14h after the final oral administration. Values represent mean ± SEM of n=2 animals per group.

## REFERENCES

1. Jara-Gutiérrez Á, Baladrón V. The Role of Prostaglandins in Different Types of Cancer. Cells. 2021;10(6):1487. doi:10.3390/cells10061487

2. Morotti M, Grimm AJ, Hope HC, et al. PGE2 inhibits TIL expansion by disrupting IL-2 signalling and mitochondrial function. Nature. 2024;629(8011):426–434. doi:10.1038/s41586-024-07352-w

3. Jin K, Qian C, Lin J, Liu B. Cyclooxygenase-2-Prostaglandin E2 pathway: A key player in tumor-associated immune cells. Front Oncol. 2023;13:1099811. doi:10.3389/fonc.2023.1099811

4. Nataraj C, Thomas DW, Tilley SL, et al. Receptors for prostaglandin E_2_ that regulate cellular immune responses in the mouse. J Clin Invest. 2001;108(8):1229–1235. doi:10.1172/JCI13640

5. Pedde AM, Kim H, Donakonda S, et al. Tissue-colonizing disseminated tumor cells secrete prostaglandin E2 to promote NK cell dysfunction and evade anti-metastatic immunity. Cell Rep. 2024;43(11):114855. doi:10.1016/j.celrep.2024.114855

6. Böttcher JP, Bonavita E, Chakravarty P, et al. NK Cells Stimulate Recruitment of cDC1 into the Tumor Microenvironment Promoting Cancer Immune Control. Cell. 2018;172(5):1022–1037.e14. doi:10.1016/j.cell.2018.01.004

7. Lacher SB, Dörr J, De Almeida GP, et al. PGE2 limits effector expansion of tumour-infiltrating stem-like CD8+ T cells. Nature. 2024;629(8011):417–425. doi:10.1038/s41586-024-07254-x

8. Obermajer N, Wong JL, Edwards RP, Odunsi K, Moysich K, Kalinski P. PGE(2)-driven induction and maintenance of cancer-associated myeloid-derived suppressor cells. Immunol Invest. 2012;41(6-7):635–657. doi:10.3109/08820139.2012.695417

9. Obermajer N, Muthuswamy R, Odunsi K, Edwards RP, Kalinski P. PGE(2)-induced CXCL12 production and CXCR4 expression controls the accumulation of human MDSCs in ovarian cancer environment. Cancer Res. 2011;71(24):7463–7470. doi:10.1158/0008-5472.CAN-11-2449

10. Bayerl F, Meiser P, Donakonda S, et al. Tumor-derived prostaglandin E2 programs cDC1 dysfunction to impair intratumoral orchestration of anti-cancer T cell responses. Immunity. 2023;56(6):1341–1358.e11. doi:10.1016/j.immuni.2023.05.011

11. Caronni N, La Terza F, Vittoria FM, et al. IL-1β+ macrophages fuel pathogenic inflammation in pancreatic cancer. Nature. 2023;623(7986):415–422. doi:10.1038/s41586-023-06685-2

12. Elewaut A, Estivill G, Bayerl F, et al. Cancer cells impair monocyte-mediated T cell stimulation to evade immunity. Nature. 2025;637(8046):716–725. doi:10.1038/s41586-024-08257-4

13. Jayaprakash V, Menezes RJ, Javle MM, et al. Regular aspirin use and esophageal cancer risk. Int J Cancer. 2006;119(1):202–207. doi:10.1002/ijc.21814

14. Lindblad M, Lagergren J, García Rodríguez LA. Nonsteroidal anti-inflammatory drugs and risk of esophageal and gastric cancer. Cancer Epidemiol Biomark Prev Publ Am Assoc Cancer Res Cosponsored Am Soc Prev Oncol. 2005;14(2):444–450. doi:10.1158/1055-9965.EPI-04-0467

15. Jacobs EJ, Rodriguez C, Mondul AM, et al. A large cohort study of aspirin and other nonsteroidal anti-inflammatory drugs and prostate cancer incidence. J Natl Cancer Inst. 2005;97(13):975–980. doi:10.1093/jnci/dji173

16. Hu T, Liu CJ, Yin X, et al. Selective COX-2 inhibitors do not increase gastrointestinal reactions after colorectal cancer surgery: a systematic review and meta-analysis. BMC Gastroenterol. 2023;23(1):281. doi:10.1186/s12876-023-02918-w

17. Ye SY, Li JY, Li TH, et al. The Efficacy and Safety of Celecoxib in Addition to Standard Cancer Therapy: A Systematic Review and Meta-Analysis of Randomized Controlled Trials. Curr Oncol Tor Ont. 2022;29(9):6137–6153. doi:10.3390/curroncol29090482

18. Martling A, Hed Myrberg I, Nilbert M, et al. Low-Dose Aspirin for PI3K-Altered Localized Colorectal Cancer. N Engl J Med. 2025;393(11):1051–1064. doi:10.1056/NEJMoa2504650

19. Wu Z, Zhang Y, Cheng Y, et al. PD-1 blockade plus COX inhibitors in dMMR metastatic colorectal cancer: Clinical, genomic, and immunologic analyses from the PCOX trial. Med N Y N. 2024;5(8):998–1015.e6. doi:10.1016/j.medj.2024.05.002

20. Menter DG, Schilsky RL, DuBois RN. Cyclooxygenase-2 and cancer treatment: understanding the risk should be worth the reward. Clin Cancer Res Off J Am Assoc Cancer Res. 2010;16(5):1384–1390. doi:10.1158/1078-0432.CCR-09-0788

21. Hong DS, Parikh A, Shapiro GI, et al. First-in-human phase I study of immunomodulatory E7046, an antagonist of PGE2-receptor E-type 4 (EP4), in patients with advanced cancers. J Immunother Cancer. 2020;8(1):e000222. doi:10.1136/jitc-2019-000222

22. Iwasa S, Koyama T, Nishino M, et al. First-in-human study of ONO-4578, an antagonist of prostaglandin E2 receptor 4, alone and with nivolumab in solid tumors. Cancer Sci. 2023;114(1):211–220. doi:10.1111/cas.15574

23. Wyrwicz L, Saunders M, Hall M, et al. AN0025, a novel antagonist of PGE2-receptor E-type 4 (EP4), in combination with total neoadjuvant treatment of advanced rectal cancer. Radiother Oncol J Eur Soc Ther Radiol Oncol. 2023;185:109669. doi:10.1016/j.radonc.2023.109669

24. Liu D, Gong J, Zhang J, et al. A phase I dose-escalation and expansion study of RMX1002, a selective E-type prostanoid receptor 4 antagonist, as monotherapy and in combination with anti-PD-1 antibody in advanced solid tumors. Invest New Drugs. 2025;43(2):250–261. doi:10.1007/s10637-025-01512-z

25. Kawazoe A, Yamaguchi K, Hamaguchi T, et al. ONO-4578 Plus Nivolumab in Unresectable Advanced or Recurrent Gastric or Gastroesophageal Junction Cancer. Cancer Sci. 2025;116(9):2523–2536. doi:10.1111/cas.70130

26. Francica BJ, Holtz A, Lopez J, et al. Dual Blockade of EP2 and EP4 Signaling is Required for Optimal Immune Activation and Antitumor Activity Against Prostaglandin-Expressing Tumors. Cancer Res Commun. 2023;3(8):1486–1500. doi:10.1158/2767-9764.CRC-23-0249

27. Punyawatthananukool S, Matsuura R, Wongchang T, et al. Prostaglandin E2-EP2/EP4 signaling induces immunosuppression in human cancer by impairing bioenergetics and ribosome biogenesis in immune cells. Nat Commun. 2024;15(1):9464. doi:10.1038/s41467-024-53706-3

28. Bonavita E, Bromley CP, Jonsson G, et al. Antagonistic Inflammatory Phenotypes Dictate Tumor Fate and Response to Immune Checkpoint Blockade. Immunity. 2020;53(6):1215–1229.e8. doi:10.1016/j.immuni.2020.10.020

29. Trotta R, Rivis S, Zhao S, et al. Activated T Cells Break Tumor Immunosuppression by Macrophage Reeducation. Cancer Discov. 2025;15(7):1410–1436. doi:10.1158/2159-8290.CD-24-0415

30. Chen Y, Perussia B, Campbell KS. Prostaglandin D2 suppresses human NK cell function via signaling through D prostanoid receptor. J Immunol Baltim Md 1950. 2007;179(5):2766–2773. doi:10.4049/jimmunol.179.5.2766

31. Corminboeuf O, Diethelm S, Zumbrunn C, et al. Design of Dual EP2/EP4 Antagonists through Scaffold Merging of Selective Inhibitors. ChemMedChem. 2024;19(2):e202300606. doi:10.1002/cmdc.202300606

32. Lyothier I, Diethelm S, Pothier J, et al. Discovery of Novel Aminopyrimidines as Selective EP2 Receptor Antagonists. ChemMedChem. 2025;20(12):e202500119. doi:10.1002/cmdc.202500119

33. Lyothier I, Diethelm S, Pothier J, et al. Discovery of ACT-1002-4271 as a Dual Prostaglandin E2 Receptor 2/Prostaglandin E2 Receptor 4 Antagonist with In Vivo Anti-Tumor Efficacy. ChemMedChem. 2025;20(12):e202500120. doi:10.1002/cmdc.202500120

34. Timmerman LM, Hensen LCM, van Eijs MJM, et al. In vitro T cell responses to PD-1 blockade are reduced by IFN-α but do not predict therapy response in melanoma patients. Cancer Immunol Immunother CII. 2024;73(9):181. doi:10.1007/s00262-024-03760-z

35. Zheng S, Wang W, Aldahdooh J, et al. SynergyFinder Plus: Toward Better Interpretation and Annotation of Drug Combination Screening Datasets. Genomics Proteomics Bioinformatics. 2022;20(3):587–596. doi:10.1016/j.gpb.2022.01.004

36. Ulahannan SV, Powderly JD, Johnson ML, et al. A phase 1 study of TPST-1495 as a single agent and in combination with pembrolizumab in patients with advanced solid tumors. J Clin Oncol. 2023;41(16_suppl):3107–3107. doi:10.1200/JCO.2023.41.16_suppl.3107

37. Davoudinasab B, Raskovalov A, Lee W, et al. Structural insights into the mechanism of activation and inhibition of the prostaglandin D2 receptor 1. Nat Commun. 2025;16(1):8944. doi:10.1038/s41467-025-64002-z

38. Zhang H, Liu Y, Liu J, et al. cAMP-PKA/EPAC signaling and cancer: the interplay in tumor microenvironment. J Hematol OncolJ Hematol Oncol. 2024;17(1):5. doi:10.1186/s13045-024-01524-x

39. Tanaka K, Hirai H, Takano S, Nakamura M, Nagata K. Effects of prostaglandin D2 on helper T cell functions. Biochem Biophys Res Commun. 2004;316(4):1009–1014. doi:10.1016/j.bbrc.2004.02.151

40. Walsh RJ, Sundar R, Lim JSJ. Immune checkpoint inhibitor combinations-current and emerging strategies. Br J Cancer. 2023;128(8):1415–1417. doi:10.1038/s41416-023-02181-6

41. Sundstrom EC, Huang X, Wiemer AJ. Anti-TIGIT therapies: a review of preclinical and clinical efficacy and mechanisms. Cancer Immunol Immunother CII. 2025;74(8):272. doi:10.1007/s00262-025-04128-7

42. af Forselles KJ, Root J, Clarke T, et al. In vitro and in vivo characterization of PF-04418948, a novel, potent and selective prostaglandin EP₂ receptor antagonist. Br J Pharmacol. 2011;164(7):1847–1856. doi:10.1111/j.1476-5381.2011.01495.x

43. Albu DI, Wang Z, Huang KC, et al. EP4 Antagonism by E7046 diminishes Myeloid immunosuppression and synergizes with Treg-reducing IL-2-Diphtheria toxin fusion protein in restoring anti-tumor immunity. Oncoimmunology. 2017;6(8):e1338239. doi:10.1080/2162402X.2017.1338239

44. Takahashi G, Asanuma F, Suzuki N, et al. Effect of the potent and selective DP1 receptor antagonist, asapiprant (S-555739), in animal models of allergic rhinitis and allergic asthma. Eur J Pharmacol. 2015;765:15–23. doi:10.1016/j.ejphar.2015.08.003

45. Sturino CF, O’Neill G, Lachance N, et al. Discovery of a potent and selective prostaglandin D2 receptor antagonist, [(3R)-4-(4-chloro-benzyl)-7-fluoro-5-(methylsulfonyl)-1,2,3,4-tetrahydrocyclopenta[b]indol-3-yl]-acetic acid (MK-0524). J Med Chem. 2007;50(4):794–806. doi:10.1021/jm0603668

46. Jumper J, Evans R, Pritzel A, et al. Highly accurate protein structure prediction with AlphaFold. Nature. 2021;596(7873):583–589. doi:10.1038/s41586-021-03819-2

47. Eastman P, Galvelis R, Peláez RP, et al. OpenMM 8: Molecular Dynamics Simulation with Machine Learning Potentials. J Phys Chem B. 2024;128(1):109–116. doi:10.1021/acs.jpcb.3c06662

48. Mobley DL, Bannan CC, Rizzi A, et al. Escaping Atom Types in Force Fields Using Direct Chemical Perception. J Chem Theory Comput. 2018;14(11):6076–6092. doi:10.1021/acs.jctc.8b00640

